# Senescent cells cluster CTCF on nuclear speckles to sustain their splicing program

**DOI:** 10.1101/2024.07.16.603680

**Authors:** Spiros Palikyras, Vassiliki Varamogiani-Mamatsi, Yajie Zhu, Shyam Ramasamy, Athanasia Mizi, Isabel Liebermann, Athanasia Stavropoulou, Ioanna Papadionysiou, Deniz Bartsch, Yulia Kargapolova, Konstantinos Sofiadis, Christoforos Nikolaou, Leo Kurian, A. Marieke Oudelaar, Mariano Barbieri, Argyris Papantonis

## Abstract

Senescence —the endpoint of replicative lifespan for normal cells— is established via a complex sequence of molecular events. One such event is the dramatic reorganization of CTCF into senescence-induced clusters (SICCs). However, the molecular determinants, genomic consequences, and functional purpose of SICCs remained unknown. Here, we combine functional assays, super-resolution imaging, and 3D genomics with computational modelling to dissect SICC emergence. We establish that the competition between CTCF-bound and non-bound loci dictates clustering propensity. Upon senescence entry, cells repurpose SRRM2 —a key component of nuclear speckles— and BANF1 —a ‘molecular glue’ for chromosomes— to cluster CTCF and rewire genome architecture. This CTCF-centric reorganization in reference to nuclear speckles functionally sustains the senescence splicing program, as SICC disruption fully reverts alternative splicing patterns. We therefore uncover a new paradigm, whereby cells translate changes in nuclear biochemistry into architectural changes directing splicing choices so as to commit to the fate of senescence.

**GRAPHICAL ABSTRACT:** 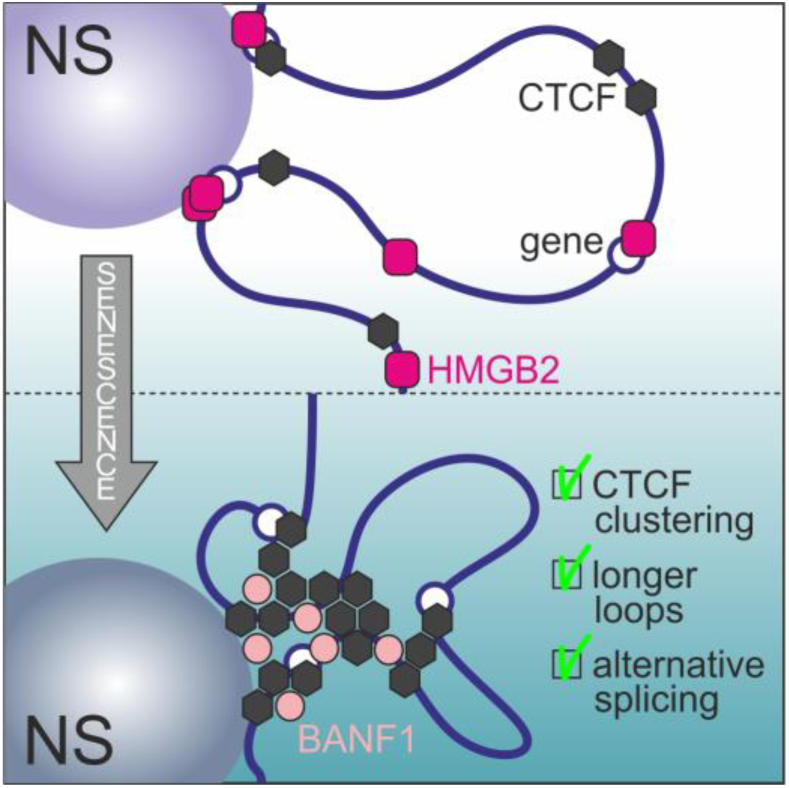

**HIGHLIGHTS:**

- HMGB2-bound loci compete with CTCF-bound ones for nuclear speckle association
- Senescent cells repurpose SRRM2 and BANF1 to cluster CTCF on speckles
- BANF1 is essential, but not sufficient for CTCF clustering
- The SRRM2 RNA-binding domain directs CTCF clustering
- SICCs rewire chromatin positioning to sustain the senescence splicing program

## INTRODUCTION

Organismal and tissue ageing is a complex phenomenon characterized by a multitude of changes to cell homeostasis. One important such change is the accumulation of senescent cells in ageing tissues^1^, clearance of which leads to life- and health-span extension in animal models^2^. Senescence can be triggered by a variety of stimuli^3^, with replicative senescence (RS) typically associated with telomere shortening over continuous cell divisions^4,5^. Alongside the irreversible, p53-mediated growth arrest that follows commitment to the state of senescence, changes in gene expression levels^6–8^, chromatin identity^6,9,10^ and genome organization ensue^8,11–16^. Many of these changes actually converge between different kinds of cells and types of senescence^7,8,13^.

As a result, cellular senescence constitutes an attractive model for studying the 3D structure-to-function relationship of human chromosomes^16,17^, in which the CCCTC-binding insulator (CTCF)^18^ holds a key role. CTCF is a conserved zinc-finger transcription factor implicated in the insulation of 3D contact domains^19–21^ and the formation of thousands of chromatin loops along chromosomes^22,23^. CTCF is also relevant in the context of senescence as, for example, the small but measurable senescence-induced decrease in CTCF levels allows activation of the *IGF2* and *INK4/ARF* loci due to reduced insulation^24,25^.

We recently described the dramatic reorganization of CTCF distribution in replicatively senescent cells into prominent senescence-induced CTCF clusters (SICCs). SICC formation is accompanied by the emergence of new, longer-range CTCF-anchored chromatin loops, which often associate with genes differentially-expressed upon senescence entry^8^. We showed that this is dependent on a key early event on the path to senescence: the nuclear loss of High mobility group-box 2 (HMGB2) proteins. HMGBs are highly abundant and ubiquitously expressed DNA-binding factors able to unwind, bend or loop DNA via interactions with their tandem HMG-box domains^26,27^. The loss of HMGB2 from human cell nuclei alone suffices for pronounced SICC formation and reduced transcriptional output genome-wide^8^.

Although SICCs represent a hallmark of senescence entry, three crucial aspects of it remain unknown. First, what are the molecular components allowing for CTCF clusters to form uniquely in the context of senescence? Second, how can the loss of the —otherwise unrelated to CTCF— HMGB2 protein trigger CTCF clustering? Third, is there a functional purpose to SICCs or are they merely a side-effect of homeostatic deregulation? Here, we address all three questions to show that SICCs form purposefully on the surface of nuclear speckles. Speckles are dynamic nucleoplasmic condensates, 20-50 nm in size, made out of poly(A)+ RNA and RNA-binding/-splicing factors^28,29^. They have been associated with storage, processing, and export of mRNAs^28,30–33^, and more recently with stress responses^34–36^. Still, the role of nuclear speckles in the transition between different cellular states, including that from homeostasis to cellular ageing, remains elusive. Here, we functionally link SICCs forming on speckles to the execution of a splicing program supporting commitment of human cells to the fate of senescence.

## RESULTS

### CTCF displays a senescence-specific protein interactome

Addressing the first question posed above, i.e., what the molecular determinants of the senescence-specific CTCF clustering are, required a cellular system that would allow us to obtain senescent cells in a defined and reproducible manner, without the heterogeneity that inevitably riddles cultures reaching replicative senescence via serial passaging. For this, we recently repurposed a small molecule inhibitor, inflachromene (ICM), initially selected for its direct targeting of HMGB1/2^37^. Treatment of primary fetal lung fibroblasts (IMR90; a popular senescence model) with ICM for 6 days produces homogeneous senescent cell populations, in which the vast majority of gene expression and phenotypic characteristics of replicative senescence manifest^38^. Importantly, ICM treatment triggers the nuclear loss of HMGB2 and prominent formation of SICCs in cells (Figures 1A and **S1A,B**), which are not a result of accumulating DNA damage (**Figure S1B**). Therefore, this system facilitates the molecular dissection of how SICCs form on an intact genome without confounding effects by population heterogeneity, paracrine signaling or asynchrony due to prolonged culturing.

**Figure 1.**
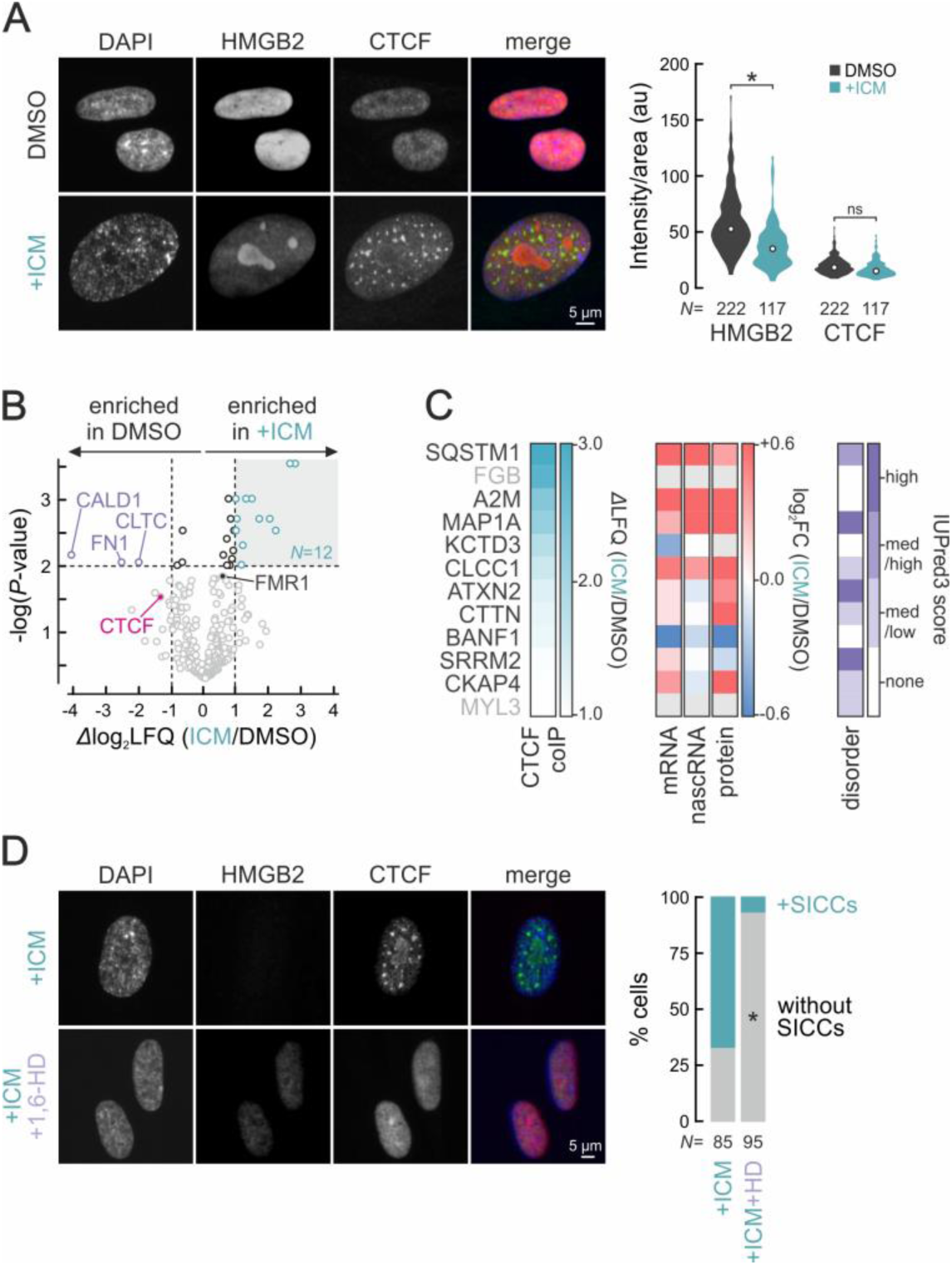
CTCF protein interactions upon senescence entry. **A**, *Left*: Representative immunofluorescence images of proliferating (DMSO) and senescent IMR90 that display SICC formation (+ICM) stained for HMGB2 and CTCF and counterstained by DAPI. *Right*: Violin plots of normalized HMGB2 and CTCF signal intensity in each condition. *N*, number of cells quantified in each condition; \**P*<0.01, two-tailed Wilcoxon-Mann-Whitney test. **B**, Volcano plot representation of proteins enriched (green) or not (purple) for interaction with CTCF based on co-immunoprecipitation-mass spectrometry data from proliferating (DMSO) and senescent IMR90 (+ICM). **C**, Heat maps showing (from left to right) *Δ*LFQ enrichments, expression levels from mRNA-seq, nascent RNA-seq and whole-cell MS/MS experiments, and predicted protein disorder based on IUPreD3 scores for the 12 significant CTCF interactors from panel B. **D**, *Left*: Representative immunofluorescence images of senescent IMR90 treated or not with 6% 1,6-hexanediol for 1 min (+1,6-HD), stained for HMGB2 and CTCF and counterstained by DAPI. *Right*: Bar plots of the percentage of cells displaying SICCs. *N*, number of cells quantified in each condition; \**P*<0.05, Fisher’s exact test.

With this system, we tested whether CTCF had a senescence-specific protein-protein interactome allowing for its clustering using co-immunoprecipitation assays coupled to mass-spectrometry in control (DMSO) and senescent IMR90 (ICM-treated). We identified 12 proteins that associated significantly more with CTCF in senescent cells (*Δ*log_2_LFQ >1, *P*-value <0.01; **Figure 1B** and **Table S1**) compared to just three that were significantly depleted. Of these 12, two —fibrinogen B chain (FGB) and myosin light chain 3 (MYL3)— were disregarded as they were not detected in our mRNA-seq, nascent RNA-seq or whole-cell proteome data from IMR90 (**Figure 1C**). The remaining 10 proteins included cytoskeletal (MAP1A, CKAP4, CTTN) and nuclear membrane components (BANF1), autophagy/mTOR signaling factors (ATXN2, SQSTM1), ion channel components (KCTD3, CLCC1), a cytokine transporter (A2M), and a key nuclear speckle protein (SRRM2). They mostly showed increased expression in senescence (7 out 10; **Figure 1C**), as well as medium to high disorder in their protein sequence (4 out 10; Figures 1C and **S1C**) as predicted by IUPred3^39^. The latter was relevant for our analysis because CTCF clusters likened condensates under the microscope (**Figure 1A**), the DNA-binding and N-terminal domains of CTCF were recently shown to coalesce *in vitro* and *in vivo*^40^, and we could readily dissolve SICCs by a short (1-min) treatment with 1,6-hexanediol (**Figure 1D**), an aliphatic alcohol interfering with weak hydrophobic interactions often required for phase separation^41^. We therefore considered these 10 factors as possible contributors to SICC formation, presumably via multivalent interactions promoting phase transitions in senescence.

### BANF1 is necessary, but not sufficient for SICC formation

To test the capacity of these factors to promote SICC formation, we shortlisted five of them. We excluded cytoskeletal components as they would be difficult to deplete from cells without major confounding defects, ion channel components due to their cell membrane anchoring (at least in non-senescent cells), and A2M as a predominantly plasma-enriched factor. We proceeded with the remaining four candidates —ATXN2, BANF1, SQSTM1, and SRRM2— for siRNA-mediated knockdowns. All four proteins had similar biophysical profiles (deduced using CIDER^42^) rendering them plausible candidates for the formation of condensates (**Figure S2A**). We also tested FMR1, an RNA-binding protein implicated in ageing-associated pathology^43^ that also displayed increased association with CTCF in senescence (**Figure 1B** and **Table S1**), shared the same biophysical properties as the other four candidates (**Figure S2A**), and carried a highly disordered C-terminus (**Figure S1C**).

Following knockdowns of *ATXN2*, *FMR1* and *SQSTM1*, and although we could efficiently deplete each one from senescent cells (by >70%; **Figure S2B**), no effect on SICC formation could be seen (**Figure S2C-E**). Therefore, we turned to BANF1 (a.k.a. BAF), which is normally part of the nuclear periphery. BANF1 functions as a ‘molecular glue’ during mitosis, wrapping anaphase chromosomes in complexes of Lamin A/C and Emerin. This holds chromosomes together while the nuclear envelope of daughter cells forms and ensures faithful genome segregation^44^. Notably, BANF1 is the only member of this group with very low sequence disorder, and the smallest in size (**Figure S1C**). BANF1 levels are reduced in senescence, but essentially all remaining protein was found in the chromatin fraction of senescent cells (Figures 1C and **2A**). BANF1 depletion from proliferating cells had no discernible effect on CTCF distribution (Figures 2B and **S3A**), but did perturb Lamin A patterns in the nuclear membrane (**Figure S3B**). BANF1 depletion from senescent cells, though, led to the near-complete abrogation of SICCs (**Figure 2B,C**). Quantification of CTCF and BANF1 levels from immunofluorescence experiments showed that SICCs cannot form once BANF1 levels drop below a critical threshold (**Figure 2D**). Since SICCs required BANF1 to form, we next asked whether increasing BANF1 levels could induce ectopic CTCF clusters. We overexpressed a BANF1-Venus fusion in proliferating IMR90 via an inducible piggybac vector^8^, but this did not produce any SICCs (**Figure 2E**). From these, we concluded that BANF1 is necessary, but not sufficient for SICC formation.

**Figure 2.**
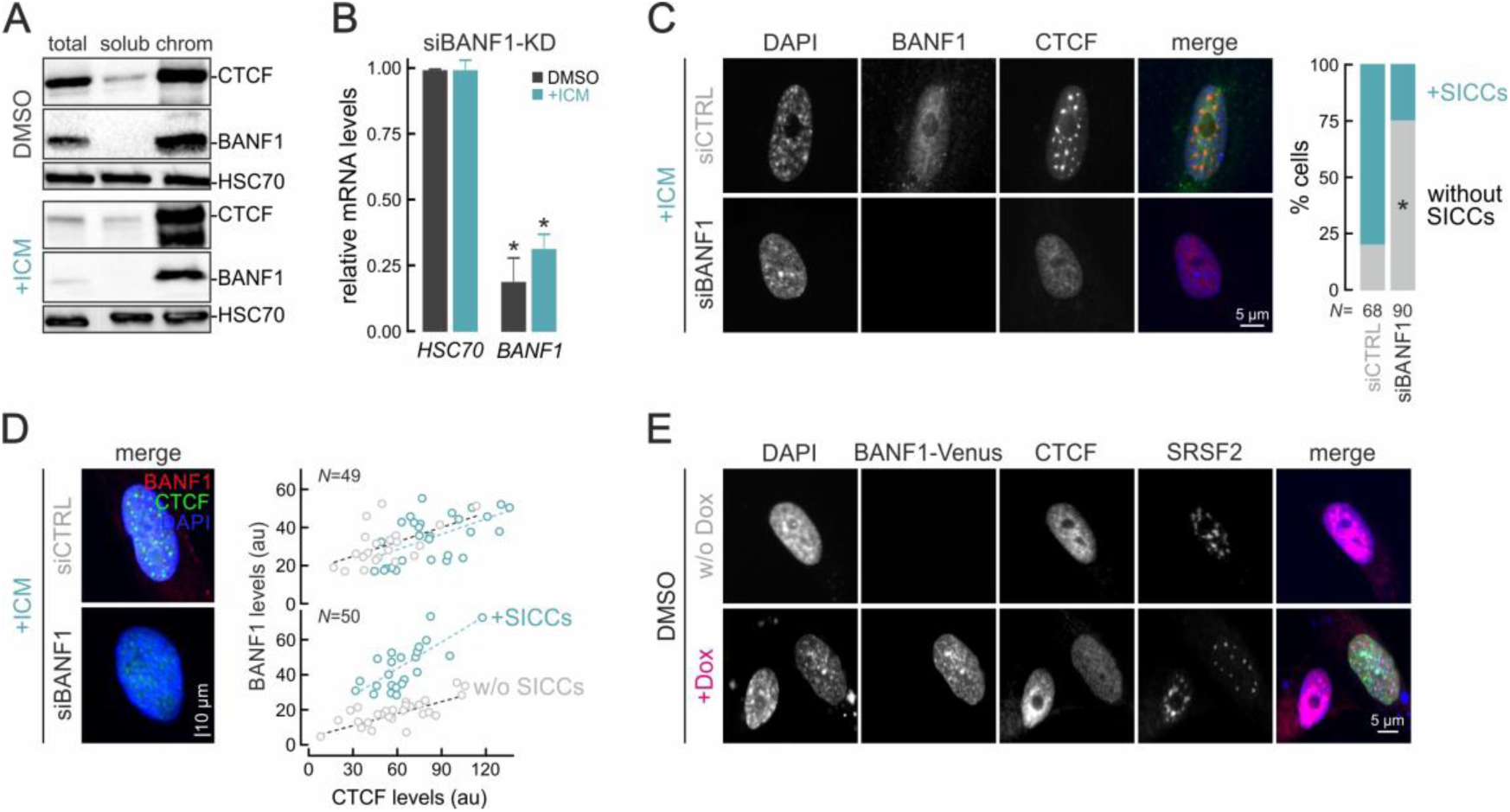
BANF1 is necessary for SICC formation. **A**, Chromatin fractionation western blots from proliferating (DMSO) and senescent IMR90 (+ICM) showing CTCF and BANF1 levels in total-cell, soluble nucleoplasm and chromatin-enriched lysates. HSC70 levels provide a loading control. **B**, Bar plots showing changes in mRNA levels upon siRNA-mediated *BANF1*-KD in proliferating (DMSO) and senescent IMR90 (+ICM) relative to *HSC70* controls. \**P*<0.01, unpaired two-tailed Student’s t-test. **C**, *Left*: Representative immunofluorescence images of senescent IMR90 treated with non-targeting (siCTRL) or *BANF1* siRNAs (siBANF1), stained for BANF1 and CTCF, and counterstained by DAPI. *Right*: Bar plots of the percentage of cells displaying SICCs. *N*, number of cells quantified in each condition; \**P*<0.05, Fisher’s exact test. **D**, *Left*: Merged immunofluorescence images like those in panel C. *Right*: Scatter plots showing single-nucleus quantifications of BANF1 and CTCF intensity from randomly-selected cells carrying (green) or not SICCs (grey). Dotted lines represent the least-squares’ fit to each data point subset. **E**, Representative immunofluorescence images of proliferating IMR90 stained for CTCF and SRSF2 and counterstained by DAPI that were treated or not with doxycycline (±Dox) to induce expression of the BANF1-Venus fusion constructs (green channel).

### Nuclear speckle association via SRRM2 directs SICC formation

The SRRM2 and SON RNA-binding proteins are the two core structural components of nuclear speckles^45^ and SRRM2 interacted strongly with CTCF in senescence (**Figure 1B,C**). We therefore first tested whether SICCs co-localize with nuclear speckles. Immunostaining of CTCF and SC-35 revealed near-perfect co-localization of SICCs and nuclear speckles (**Figure 3A**). In fact, we manipulated the shape and numbers of nuclear speckles using pharmacological inhibitors^46^, and SICC reshaped and redistributed following how speckles changed: tautomycetin disperses nuclear speckles^47^ and SICCs dispersed accordingly upon treatment (**Figure S4A**), while cantharidin forces speckle fusion^48^ and SICCs then also fused together (**Figure S4B**).

**Figure 3.**
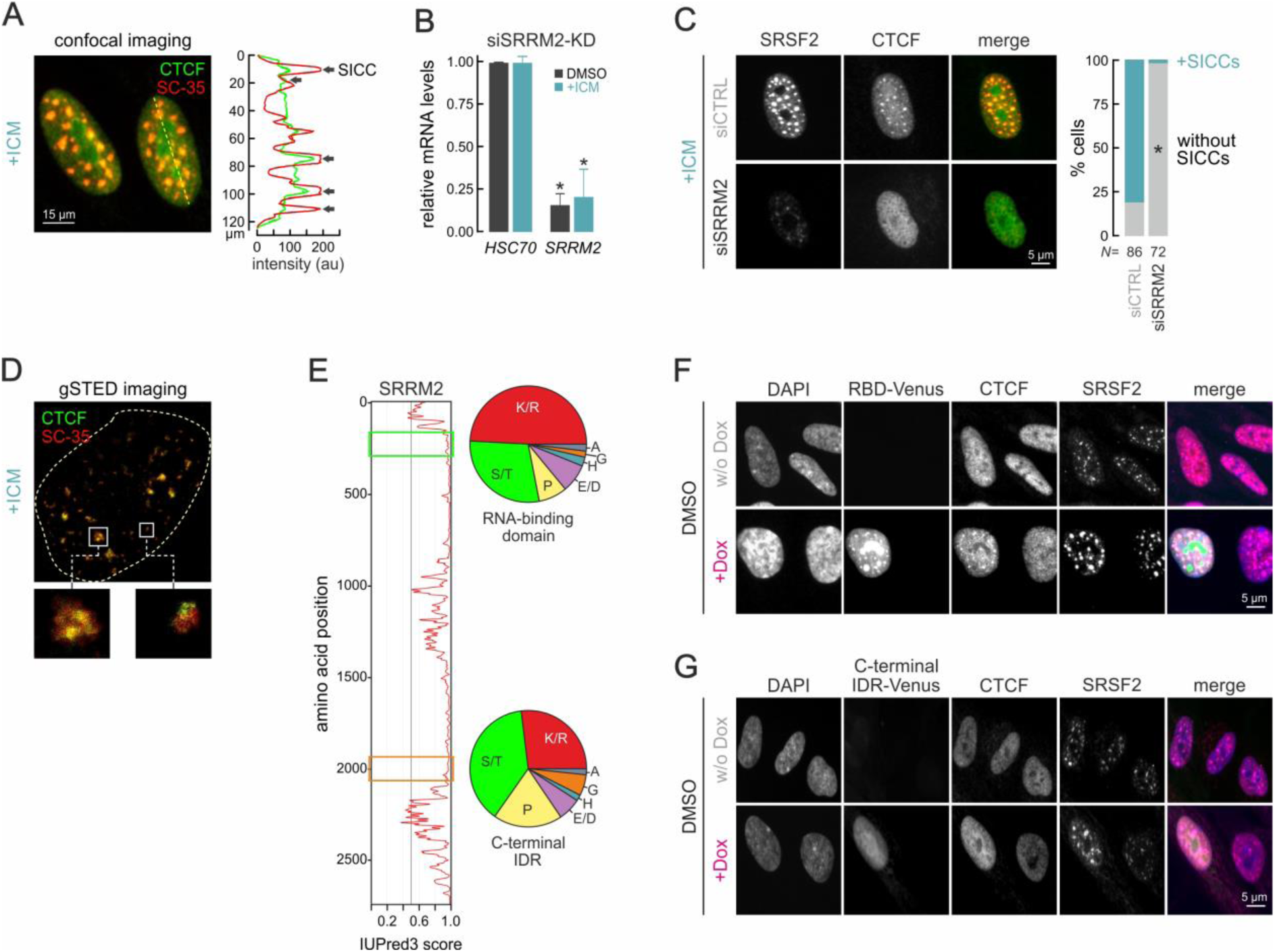
SRRM2 is necessary and sufficient for SICC formation. **A**, *Left*: Representative immunofluorescence image from senescent IMR90 (+ICM) stained for CTCF and the SC-35 speckle marker. HSC70 levels provide a loading control. *Right*: Co-localization plot of CTCF and SC-35 fluorescence intensity signal (arrows) in the area under the dotted line on the confocal image. **B**, Bar plots showing changes in mRNA levels upon siRNA-mediated *SRRM2*-KD in proliferating (DMSO) and senescent IMR90 (+ICM) relative to *HSC70* controls. \**P*<0.01, unpaired two-tailed Student’s t-test. **C**, *Left*: Representative immunofluorescence images of senescent IMR90 stained for SRSF2 and CTCF and counterstained by DAPI treated with non-targeting (siCTRL) or *SRRM2* siRNAs (siSRRM2). *Right*: Bar plots of the percent of cells displaying SICCs. *N*, the number of cells quantified in each condition; \**P*<0.05, Fisher’s exact test. **D**, Super-resolution imaging of a senescent IMR90 nucleus (dotted line) stained for CTCF and SC-35. Two characteristic SICCs overlapping nuclear speckles (rectangles) are magnified. **E**, Line plot showing the predicted IUPred3 disorder score along SRRM2. The 68 aa-long segments of the protein’s RNA-binding domain (RBD) and C-terminal intrinsically disordered region (IDR) are marked (rectangles) and their amino acid compositions shown (pie charts). **F**, Representative immunofluorescence images of proliferating IMR90 stained for CTCF and SRSF2 and counterstained by DAPI that were treated or not with doxycycline (±Dox) to induce SRRM2 RBD-Venus expression of the fusion constructs (green channel). **G**, As in panel F, but to induce SRRM2 IDR-Venus expression (green channel).

We next knocked-down *SRRM2* in senescent IMR90 and completely disrupted SICCs (**Figure 3B,C**). This occurred despite speckles not fully collapsing (**Figure S5A**) as would be expected^45^. Dependence of SICC formation on SRRM2 also held true in proliferating cells depleted of HMGB2 to induce CTCF clustering (**Figure S5B**). Given this data, we revisited our 1,6-hexanediol experiments that disrupted SICCs (**Figure 1D**) and found that speckles did not dissolve during the treatment (**Figure S5C**). The CTCF-SRRM2 interaction is therefore necessary for SICC formation, not just speckle integrity.

RNA species often promote condensate formation, including that of nuclear speckles^49^. In addition, CTCF was reported to require interaction with RNA in order to establish a subset of loops in the genome^50,51^, although the specific nature of these interactions is now disputed^52^. We therefore decided to test the contribution of RNA to SICCs. We first treated cells with ammonium acetate, known to disrupt RNA-protein condensates (e.g., stress granules^53^). Applying ammonium acetate at a concentration that does not disrupt nuclear speckles had no effect on SICCs (**Figure S6A**). We next treated cells with RNase A, before fixing and immunostaining them for CTCF and speckle markers. Although speckles displayed some dispersal, SICCs were not compromised by the degradation of cellular RNA (**Figure S6B**). This was consistent with co-IP experiments showing that interaction between CTCF and speckles was markedly strengthened upon RNA, but abolished by DNA enzymatic degradation in senescent cells (**Figure S6C**). We also performed eCLIP^54^ targeting CTCF in both proliferating and senescent IMR90. In line with recent biochemical evidence^52^, we found very limited association of CTCF with RNA in proliferating cells. However, in senescence and upon SICC formation on speckles, we mapped 559 eCLIP peaks in >250 mRNAs (**Figure S6D,E**). 55% of these peaks mapped to the 5’ end of target transcripts, and these CTCF targets were linked to GO terms like ‘RNAPII regulation’ and ‘aging’ (**Figure S6F,G**). Then, if anything, RNA appears refractory to the CTCF-speckle co-association. This agrees with gSTED super-resolution imaging of SICCs and nuclear speckles in senescent cells, where the two entities did not appear to mix, but SICCS rather formed on the surface of speckles (**Figure 3D**). Last, given that RNA was proposed to mediate both CTCF-CTCF and CTCF-DNA interactions^50,51^, we decided to manipulate in-cell Zn^+2^ availability to interfere with CTCF binding to chromatin. The Zn^+2^ chelator TPA, shown to increase the fraction of unbound CTCF^55^, did not affect SICC formation. However, increasing Zn^+2^ titers, shown to enhance CTCF chromatin association^55^, reduced SICC formation significantly (**Figure S6H**). Thus, unbound CTCF is likely recruited to CTCF clusters and contributes to their emergence.

Finally, we tested whether increasing SRRM2 levels would enforce ectopic CTCF clustering. For this, we cloned a 63-aa portion of the SRRM2 N-terminal IDR encompassing its RNA-binding domain in the same piggybac vector as above (**Figure 3E**). Overexpression of this SRRM2 peptide in proliferating IMR90 sufficed for the specific induction of CTCF clustering at nuclear speckles (and not in nucleoli, where the fusion peptide also accumulated; **Figure 3F**). This highly selective effect held also true in cells, senescent and proliferating alike, from which we had first depleted SRRM2 (**Figure S7A**). To assess if this could be achieved by any IDR, we overexpressed a 63 aa-long portion of the SRRM2 C-terminal IDR that was of relatively similar amino acid composition as its RNA-binding domain (Figures 3E, **S2A** and **S7B**). However, overexpression of this disordered peptide could not induce SICCs (**Figure 3G**). These results showed that SRRM2 (and, in particular, its RNA-binding domain) is both necessary and sufficient for SICC formation. Together with our data on BANF1, we addressed our first question and identified two factors that are indispensable drivers of CTCF clustering on nuclear speckles upon entry into senescence.

### Competition for nuclear speckle association underlies CTCF clustering

We next addressed the second question we posed at the beginning, i.e., how the nuclear loss of HMGB2 triggers CTCF clustering. Previously, we were able to prevent SICC formation in senescent nuclei by re-introducing HMGB2^8^. Here, we reiterated this, while also testing the effects of two HMGB2 subdomains: its 71 aa-long DNA-binding ‘box A’ and its disordered and highly acidic 48-aa ‘tail’ (Figures 4A and **S2A**). As shown before, overexpression of full length HMGB2-Venus in senescent cells resulted in the strong reduction of cells with SICCs compared to Venus alone (**Figure 4B**). This effect was faithfully recapitulated upon overexpression of just the HMGB2 acidic tail, but not of its DNA-binding domain (**Figure 4C**), suggesting that its disordered tail alone interferes with the CTCF-BANF1-SRRM2 interaction and deter clustering.

**Figure 4.**
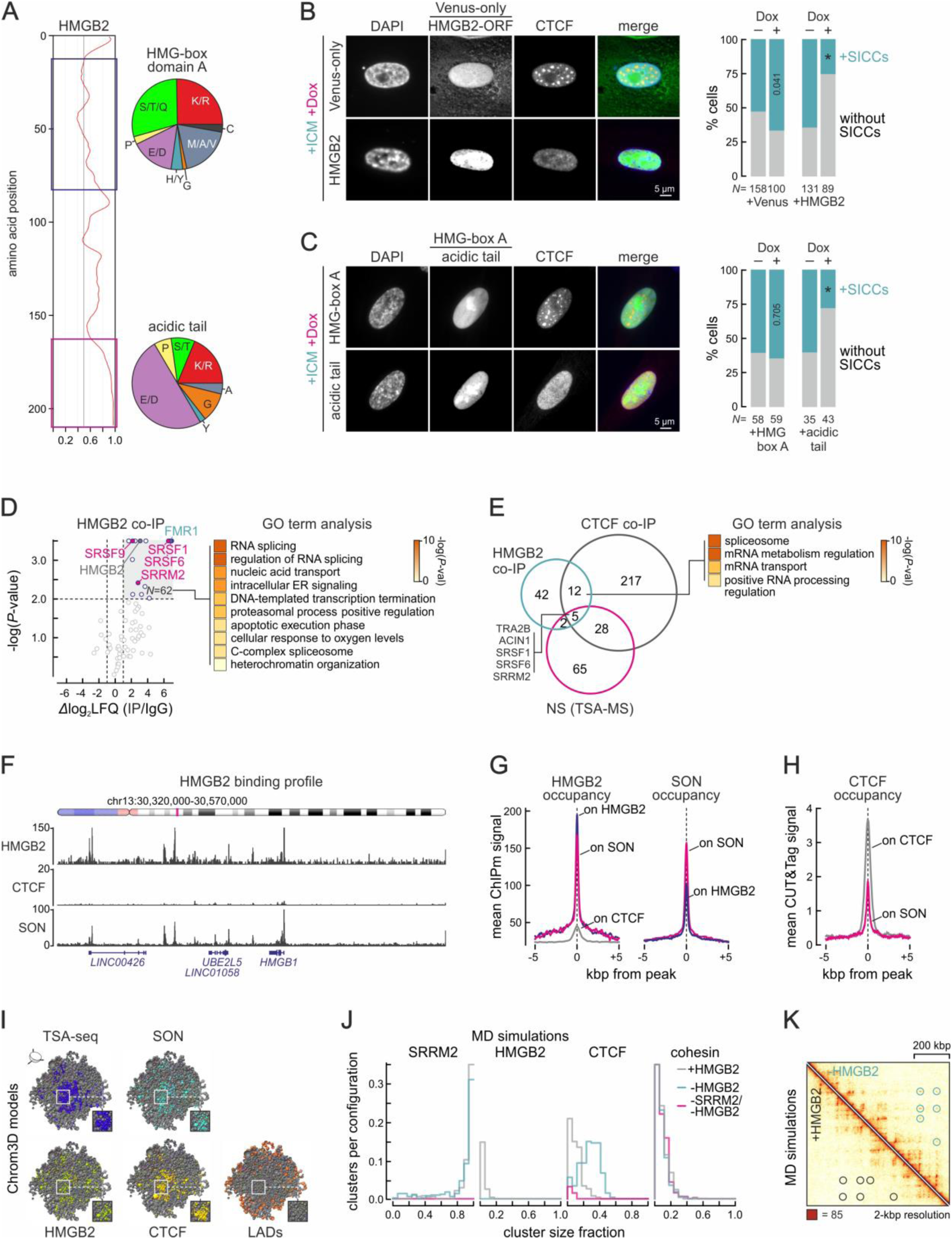
HMGB2 competes with CTCF for speckle association. **A,** Line plot showing the predicted IUPred3 disorder score along HMGB2. The protein’s HMG-box A DNA-binding domain and C-terminal acidic tail are marked (rectangles) and their amino acid compositions shown (pie charts). **B,** *Left*: Representative immunofluorescence images of senescent IMR90 (+ICM) stained for CTCF and counterstained by DAPI that were treated or not with doxycycline (±Dox) to induce expression of the HMG-box A fusion constructs (green channel). *Right*: Bar plots of the percentage of cells displaying SICCs. *N*, the number of cells quantified in each condition; \**P*<0.05, Fisher’s exact test. **C,** As in panel B, but treated or not with doxycycline (±Dox) to induce expression of the HMGB2 C-terminal acidic tail-Venus fusion constructs (green channel). **D,** *Left*: Volcano plot representation of proteins enriched (green) or not (purple) for interaction with HMGB2 based on co-immunoprecipitation-mass spectrometry data in proliferating IMR90 (DMSO) over IgG controls. *Right*: Heat map showing the top ten most enriched GO terms/pathways associated with the 62 significant HMGB2 protein interactors. **E,** *Left*: Venn diagram showing the overlap between all HMGB2-(green circle) and CTCF-interacting proteins (grey circle) from co-immunoprecipitation-mass spectrometry data in proliferating and senescent IMR90, respectively, with the top 100 nuclear speckle (NS) components based on TSA-MS data^56^. *Right*: Heat map showing the GO terms/pathways associated with the 17 shared HMGB2-CTCF interactors. **F,** Representative genome browser view of a 250 kbp-long region on chr13 (ideogram) showing HMGB2 and SON ChIPmentation and CTCF CUT&Tag binding profiles from proliferating IMR90. **G,** Line plots showing mean HMGB2 (left) and SON ChIPmentation signal (right) in the 10 kbp around HMGB2 (blue), SON (magenta) or CTCF peaks (grey) from proliferating IMR90. **H,** As in panel G, but for CTCF CUT&Tag signal in the 10 kbp around SON (magenta) or CTCF peaks (grey). **I,** Sections of the Chrom3D diploid genome model from proliferating IMR90 showing the top 20% of TADs (beads) carrying most TSA-seq (blue), SON (light blue), HMGB2 (green), CTCF (yellow) or LAD signal (brown). Magnifications of HMGB2-SON overlapping regions are provided (rectangles). **J,** Plots showing the number of SRRM2, HMGB2, CTCF and cohesin clusters and their sizes in MD simulations with (grey line) and without HMGB2 (green line) or without HMGB2 and SRRM2 (magenta dotted line). **K,** Contact matrix of a 2-Mbp genomic region from chr9 deduced from MD simulations with (bottom) or without HMGB2 (top). Long-range loops forming in the absence of HMGB2 are demarcated (circles).

As HMGB2 and CTCF do not directly associate with one another^8^, we reanalyzed our HMGB2 co-IP mass-spec data looking for nuclear speckle components and other shared interactors. In proliferating cells, HMGB2 co-purifies with 62 proteins significantly enriched over background (*Δ*log_2_LFQ >1, *P*-value <0.01; **Figure 4D**). These could be linked with higher confidence to RNA-than to DNA-templated functions (**Figure 4D**). Of the 62 interactors, only two were shared with the senescent-specific CTCF interactome —FMR1 and SRRM2. However, if we consider the full list of CTCF interactors (*N*=234; **Table S1**), this number grows to 17 (i.e., >27% of the whole HMGB2 co-IP list). These 17 shared proteins are, again, almost exclusively associated with RNA splicing (**Figure 4E**). Moreover, HMGB2 interacts with seven *bona fide* components of nuclear speckles and CTCF with 33 (using the top 100 speckle-enriched proteins from TSA-MS experiments^56^ as a reference), the two of them sharing five key components including SRRM2 (**Figure 4E**). Therefore, both CTCF and HMGB2 have an inherent affinity for nuclear speckle association.

To further assess this, we generated chromatin binding profiles for HMGB2 and SON, the other key structural component of nuclear speckles together with SRRM2, which was shown to associate with chromatin^57^. We applied a sensitive ChIPmentation approach^58^ to proliferating cells, which allowed us to map >5300 and >22,000 high confidence HMGB2 and SON binding peaks, respectively (**Figure 4F**). This represented a substantial improvement compared to the ∼1100 HMGB2 peaks we were able to map before^8^, as well as to the very broad SON domains mapped using GO-CaRT (a modified CUT&RUN approach)^57^. Analysis of HMGB2 binding revealed that >75% of its peaks overlapped active genes (with ∼50% overlapping gene promoters; **Figure S8A**). Of the 2654 HMGB2-bound genes, ∼10% were up- and ∼20% were down-regulated upon senescence (|log_2_FC|>0.6; **Figure S8B**). Gene set enrichment analysis (GSEA) showed that these differentially-expressed and HMGB2-bound genes were linked to the suppression of cell cycle progression and to proinflammatory signaling, but also to RNA splicing and processing control (**Figure S8C**). Also, as we showed before^8^, HMGB2 binding was enriched at the boundaries of TADs (called at 20-kbp resolution using Micro-C data from control and ICM-treated cells^38^; **Figure S8D**). At the same time, 88% of HMGB2 and >75% of SON peaks reassuringly resided in the A1 subcompartment (**Figure S8E**) known to associate with nuclear speckles (e.g., based on TSA-seq data^59^). Notably, HMGB2 and SON overlapped at thousands of positions genome-wide (i.e., more than half of HMGB2 peaks overlap 80% of a SON peak), which was not the case for HMGB2 and CTCF (i.e., <1/8 of HMGB2 peaks overlap 80% of a CTCF peak; **Figure 4G**). On the other hand, SON and CTCF did co-occupy a large number of positions genome-wide (i.e., >1/4 of CTCF peaks overlap 80% of a SON peak; **Figure 4H**), with only 662 positions were bound by all three factors.

Given the binding of HMGB2, SON, and CTCF to the A1 subcompartment (**Figure S8E**), we used TADs deduced from our IMR90 Micro-C data and coordinates of constitutive lamina-associated domains (LADs)^60^ as input in Chrom3D^61^ to model coarse-grained whole-genome 3D organization and HMGB2/SON/CTCF distribution. Chrom3D models chromosomes as polymers of beads, each corresponding to a TAD; statistical positioning of each bead in a diploid genome is calculated on the basis of inter- and intrachromosomal Micro-C interaction signal between TADs, with LAD coordinates forcing the relevant TADs towards the nuclear periphery. The resulting models indeed showed LADs distributed peripherally, while IMR90 TSA-seq signal^59^ (the top 20%) predominantly occupied the core of the model (**Figure 4I**). Interestingly, when we highlighted the top 20% of TADs carrying most HMGB2 or SON ChIPmentation signal, we found remarkable overlap with TSA-seq TADs (i.e., ∼60% of HMGB2 and >50% of SON TADs were TSA-seq–positive) as well as with each other (i.e., ∼75% of HMGB2 TADs overlapped SON ones; **Figure 4I**). This held true for TADs carrying the top 20% of CTCF signal (i.e., >52% and ∼68% of CTCF beads overlapped SON and TSA-seq ones, respectively; **Figure 4I**), further supporting the notion that nuclear speckles lie in spatial proximity to active, HMGB2-rich chromatin domains.

Finally, we used all this information as input in molecular dynamics simulations via which we wanted to test how CTCF-bound chromatin and CTCF itself reorganize in relation to speckles following HMGB2 depletion from the nuclear milieu. We expanded our previous framework for modeling 3D chromatin conformation^62^ by adding the following features deduced from our experiments here: (i) HMGB2 interacts with nuclear speckles (**Figure 4D,G**); (ii) HMGB2 loss reduces overall transcriptional output^8^; (iii) in the absence of HMGB2, CTCF displays increased interaction with speckles (Figures 1B and **3C-G**); (iv) chromatin-bound CTCF is needed to nucleate clustering at speckles with RNA counteracting this (**Figure S6C**); and (v) the unbound CTCF fraction contributes to SICCs (**Figure S6H**). We implemented these features as weak forces in our simulations of a 3-Mbp stretch of chromatin with features copied from chr9 (see Methods). HMGB2 removal from these simulations (to mimic senescence entry) led to the formation of more and markedly larger clusters of CTCF (Figures 4J and **S8G**). Clustering on speckles was evidenced by the increased contact probability between CTCF and SRRM2 (**Figures S8H**) the latter inherently phase-separating into speckle-like structures (Figures 4J and **S8G**). Reassuringly, CTCF clustering in our models did not also result in the clustering of loop-extruding cohesin (**Figure 4J**) much like what was observed experimentally^8^. On the other hand, removal of SRRM2 from the system precluded CTCF clustering again without affecting cohesin (**Figure 4J**). Last, our MD simulations, when visualized as contact matrices, predicted formation of longer-range, nested CTCF loops upon removal of HMGB2 (**Figure 4K**). Together, our experiments and simulations describe an antagonistic relationship between HMGB2 and CTCF in homeostasis, which tilts in favour of CTCF upon the loss of HMGB2 from senescent nuclei.

### CTCF clustering drives genome reorganization in senescence

To begin addressing the third question we posed above, i.e., what might the functional purpose of SICCs be, we mapped the binding profiles of SON (via ChIPmentation) and CTCF (via CUT&Tag) in senescence. Comparison of SON profiles from proliferating and ICM-treated cells revealed binding to essentially the same positions, albeit with diminished intensity in senescence (**Figures 5A,B** and S9A). This translated into a 4-fold reduction in the number of peaks called from >22,000 in proliferating to 5435 in senescent cells, most of which mapped to promoters and genes bodies in the A1 subcompartment (**Figure S8A,E**). CTCF was also found to bind essentially the same positions genome-wide in senescence with a moderate decrease in signal (**Figure 5A,C** and S9A). However, this translated into a ∼30% increase in peaks (filtered for the presence of a consensus CTCF motif) from 6565 to >8500 upon ICM treatment as previous weak peaks gained in signal. Still, negligible change in genomic or compartmental distribution was observed for either protein (**Figure S8A,E**). It then follows that SON- and CTCF-bound positions remained strongly overlapping also in senescent cells, where HMGB2 is no longer present (Figures 5A and **S9A**). In fact, the 435 promoters bound by both factors belonged to genes associated with hallmark GO terms for senescence entry like ‘RNP biogenesis’, ‘mitotic cell cycle’, ‘cellular stress’ and ‘DNA damage response’, the ‘p53 pathway’, ‘chromosome organization’, and even ‘membraneless organelle assembly’ (**Figure 5D**), although only ∼10% of these genes changed their expression levels significantly (i.e., |log_2_FC|>0.6) upon senescence entry.

**Figure 5.**
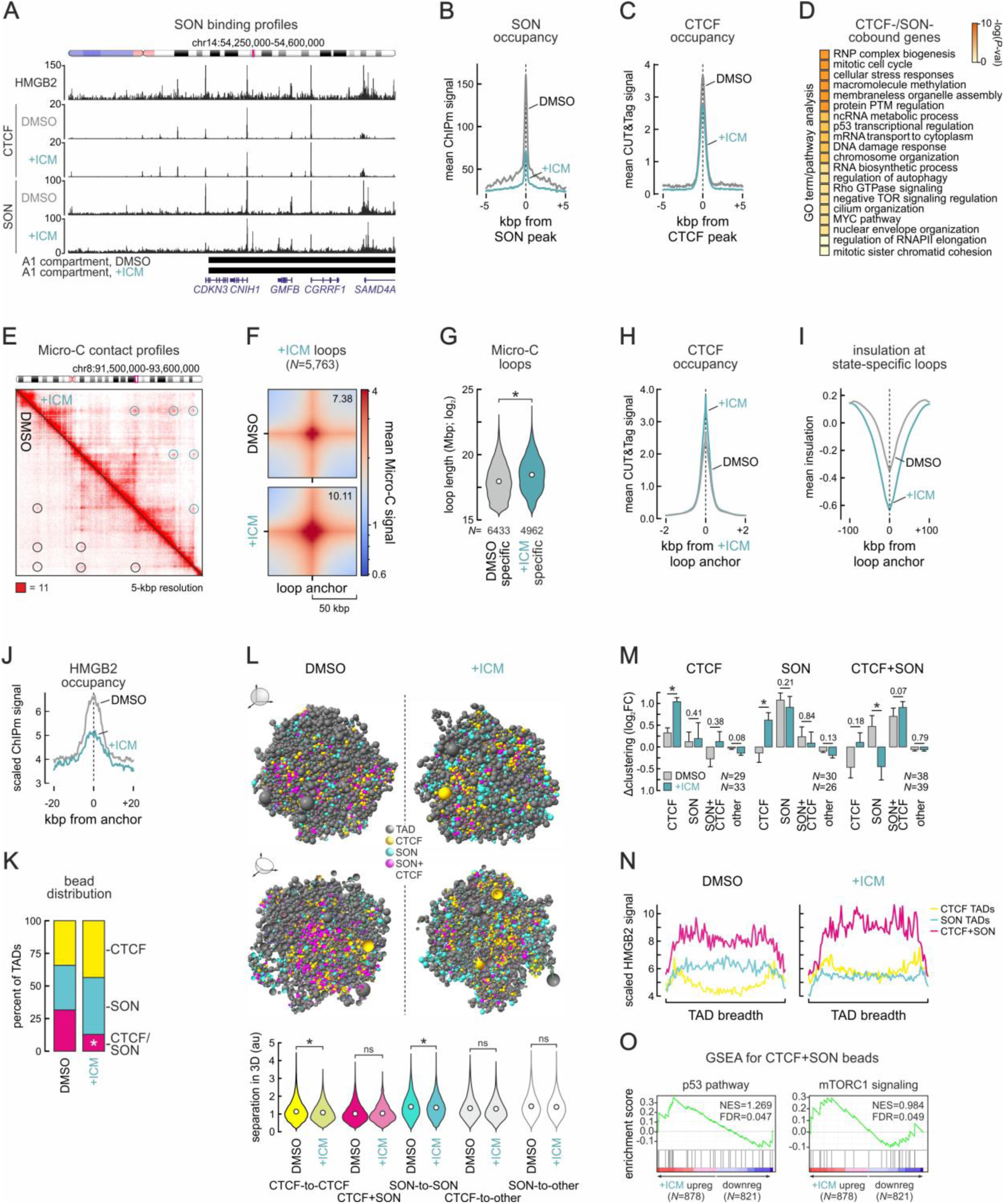
3D genome reorganization in respect to nuclear speckles and SICCs. **A**, Representative genome browser view of HMGB2 and SON ChIPmentation data aligned to CTCF CUT&Tag profiles from proliferating (DMSO) and senescent IMR90 (+ICM) in a 350-kbp region on chr14. **B**, Mean SON occupancy from proliferating (grey) and senescent IMR90 (green) in the 10 kbp around SON peaks. **C**, As in panel B, but for CTCF occupancy in the 10 kbp around CTCF peaks. **D**, Heat map showing the top most enriched GO terms/pathways associated with SON-/CTCF-cobound genes. **E**, Contact matrix showing Micro-C interactions from proliferating (DMSO) and senescent IMR90 (+ICM) in a 2.1-Mbp region from chr8. CTCF loops are indicated (black/green circles). **F**, APA plots showing mean Micro-C signal strength in proliferating (DMSO) and senescent cells (+ICM) for all loops from senescent cells. **G**, Box plots showing length distribution of loops specific to proliferating (DMSO) and senescent cells (+ICM). \**P*<0.01; Wilcoxon-Mann-Whitney test. **H**, As in panel B, but for CTCF occupancy in the 4 kbp around senescence loop anchors. **I**, Plots showing mean insulation scores from proliferating (DMSO) and senescent Micro-C (+ICM) in the 200 kbp around senescence loop anchors. **J**, As in panel B, but for HMGB2 occupancy in the 40 kbp around senescence loop anchors. **K**, Bar plots showing the percent of TADs in Chrom3D models marked by high CTCF (yellow), SON (light blue) or CTCF+SON signal (magenta). \**P*<0.01; Fisher’s exact test. **L**, *Top*: Renderings of Chrom3D diploid genome models from proliferating (DMSO) and senescent IMR90 (+ICM) showing TADs (beads) that carry the top 20% of CTCF (yellow), SON (light blue), or CTCF+SON signal (magenta). *Bottom*: Violin plots showing mean pairwise 3D separation of the indicated types of beads in each model. \**P*<0.01; Wilcoxon-Mann-Whitney test. **M**, Bar plots showing clustering (±SEM) of CTCF (left), SON (middle) or CTCF+SON TADs (right) relative to randomized controls in DMSO-(grey) or ICM-treated cells (green). \**P*<0.05; unpaired two-tailed Student’s t-test. The number (*N*) of clusters deduced in each case is indicated. **N**, Plots showing scaled HMGB2 occupancy within CTCF-(yellow), SON-(light blue) and CTCF+SON-enriched TADs (magenta) from the proliferating (DMSO) and senescent Chrom3D models (+ICM) from panel L. **O**, GSEA analysis of differentially expressed genes in the CTCF+SON-enriched TADs from panel L.

Since SON and CTCF binding along the 1D genome did not change that significantly, CTCF should reorganize the senescent genome mainly in 3D. To assess this, we used 5-kbp-resolution Micro-C data to call 7234 and 5763 chromatin loops in proliferating and senescent IMR90, respectively. Despite this 20% reduction in loops, we could record the emergence of multiple new, stronger, and usually nested loops in senescence (**Figures 5E,F** and S9B). These new loops showed increased lengths compared to proliferation-specific loops (**Figure 5G**) corroborating our previous observations by HiChIP^8^. Senescence-specific loops showed increased CTCF occupancy at their anchors (despite a general decreased in CTCF binding strength genome-wide; **Figure 5H**), as well as significantly stronger insulation (Figures 5I and **S9C**). Interestingly, >25% of HMGB2 peaks in proliferating cells fell within 10 kbp of a loop anchor (overlapping a total of 1422 anchors), whilst <16% of HMGB2 peaks fell within the same window around senescence loop anchors (overlapping 915 anchors; **Figure 5J**). Moreover, 84% of HMGB2-marked loop anchors in proliferating cells were also marked by a SON peak in contrast to <28% in senescence, meaning that loop reorganization correlated with the loss of HMGB2 from SON-bound anchors. In fact, loops with one CTCF- and one HMGB2-bound anchor were rewired in senescence, while loops with both anchors bound by HMGB2 were lost and the smallest in size (**Figure S9D-F**) explaining how HMGB2 loss fuels 3D genome reorganization upon senescence entry.

To visualize the spatial clustering of chromatin domains relative to CTCF and nuclear speckles in the senescent genome, we again turned to Chrom3D. We initially modeled the distribution of the top 20% of TADs from proliferating and ICM-treated cells that carried most CTCF signal. This revealed spatial co-association of CTCF-rich TADs that clustered significantly more upon ICM treatment (**Figure S9G,H**). We next marked the top 20% of TADs in our proliferating and senescent whole-genome models according to whether they carried CTCF or SON signal. This revealed ∼31% of TADs marked by both CTCF and SON in proliferating data. This ratio changed significantly in senescence with CTCF+SON-high TADs reduced to <13% (Figures 5K and **S9I**). Analysis of the rendered models revealed increased homotypic clustering of CTCF- and of SON-marked TADs, whereas CTCF/SON ones did not come into further 3D proximity (**Figure 5L**). Unbiased clustering analysis confirmed changes in clustering in senescence: CTCF-marked TADs associate significantly more with one another in senescence, as well as with TADs marked by both CTCF and SON (**Figure 5M**). At the same time, SON-marked TADs cluster closer to CTCF ones, while CTCF+SON TADs show more association with CTCF TADs and significantly less with SON ones (**Figure 5M**). Strikingly, double-marked TADs carried significantly more HMGB2 signal prior to senescence entry (**Figure 5N**). This indicates that CTCF-HMGB2 competition for speckle association also manifests here, despite not being presumed in our modeling parameters. All effects held also true in Chrom3D models of individual chromosomes from these two cellular states (**Figure S9J-L**). Finally, using a list of 1699 differentially expressed genes (|log_2_FC|>0.6) embedded in these CTCF+SON TADs in GSEA revealed significant association with p53 activation and mTORC1 signaling cascades (**Figure 5O**), known to control cell cycle progression and cell growth^63^. This suggests that 3D reorganization of senescent chromosomes in respect to nuclear speckles could support the gene expression program underlying senescence entry.

### SICCs sustain the senescence splicing program

Despite the findings above, the functional role of SICCs during senescence commitment remained only partially addressed. We previously showed by a combination of nascent RNA-seq, mRNA-seq, Ribo-seq and whole-cell proteomics that both replicative and ICM-induced senescence are driven predominantly by changes at the level of transcription^14,38^. Given that nuclear speckles are now shown to contribute to splicing regulation^33,45,64,97^, we hypothesized that alternative splicing (AS) might be an underappreciated driver of senescence entry.

To evaluate this, we first used *IsoformSwitch*^65^ to gauge AS consequences on mRNA isoforms and their coding potential. We found ~1000 changes in isoforms induced by ICM and senescence entry, predominated by changes involving alternative transcription start (TS) and end (TE) sites (**Figure 6A** and **Table S2**). ICM-induced isoform changes produced mature mRNAs with particular functional consequences, including loss or gain of functional domains and signal peptides, changes in ORF length, and NMD-sensitive or non-coding isoforms from many genes (Figures 6B and **S10A**). A subset (∼50%) of these effects was phenocopied upon *HMGB2* knockdown (**Figures 6A,B** and S10B) and concerned processes central to senescence entry (**Figure S10E**). Similar analyses of *SRRM2*- and *BANF1*-KD data revealed that many ICM-induced changes were reverted or diminished under conditions where SICCs cannot form (**Figures 6A,B** and **S10C,D**).

**Figure 6.**
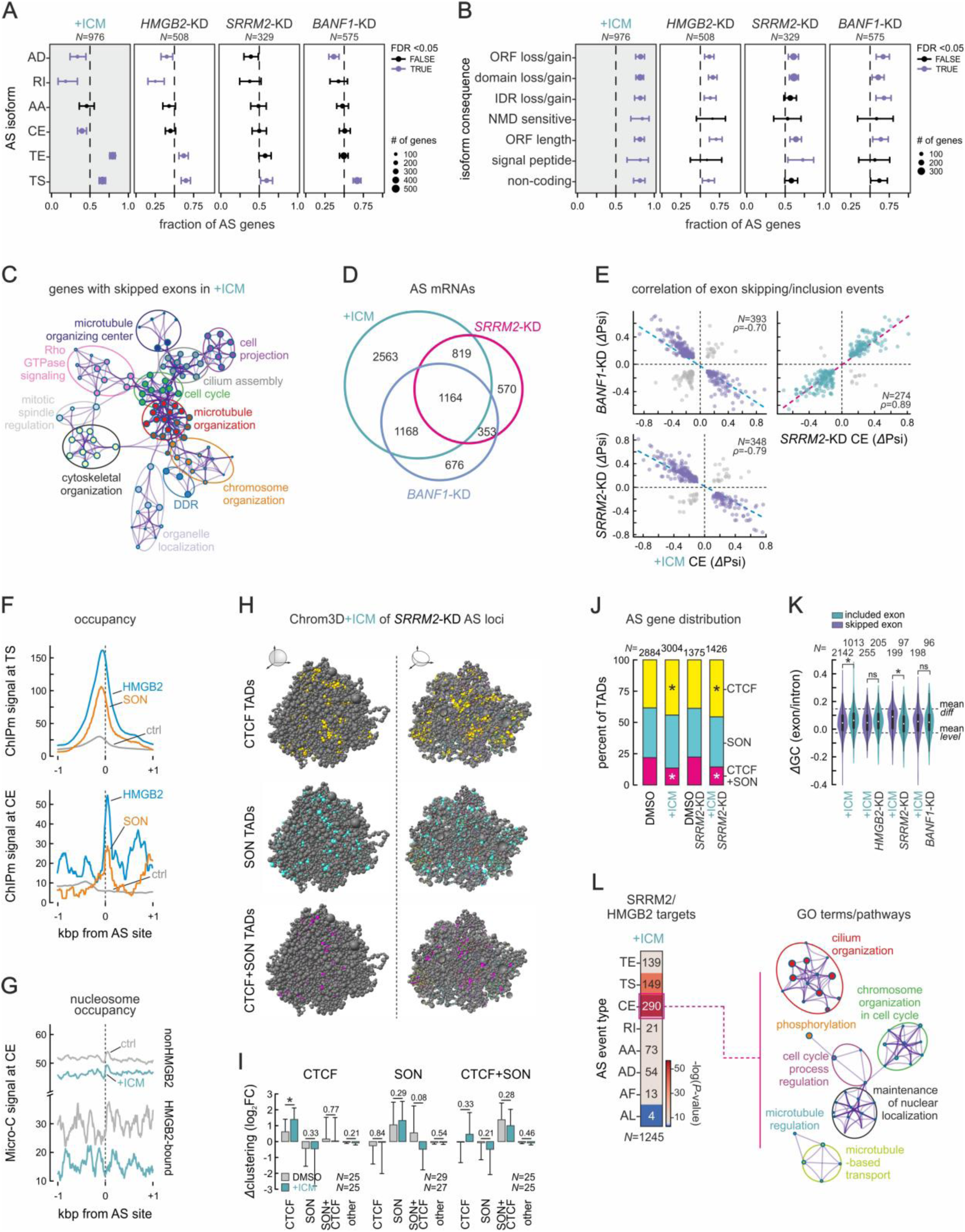
Splicing changes related to SICC formation in senescence. **A**, Plots showing the fraction of alternatively spliced (AS) genes associated with alternative donor site (AD) usage, intron retention (RI), alternative acceptor (AA) site usage, core exon skipping or inclusion (CE), and alternative transcription end (TE) or start site (TS) usage induced by senescence (+ICM) or by the knockdown (KD) of *SRRM2*, *HMGB2*, and *BANF1*. The number of AS isoforms (*N*) per each dataset is shown; AS events with an FDR<0.05 are indicated (in purple). **B**, As in panel A, but for the fraction of alternatively spliced (AS) genes associated with the indicated consequences on the encoded mRNA isoforms. **C**, Network graph showing GO terms/pathways significantly associated with CE skipped in senescence. **D**, Venn diagram showing the overlap of AS genes between senescence (+ICM) and *SRRM2*- or *BANF1*-KD. **E**, Scatter plots showing correlation of *Δ*Psi values of CE events shared between senescence (+ICM) and *SRRM2*- or *BANF1*-KD. The number of CE events (*N*) and correlation coefficients (ρ) are indicated. **F**, Line plots showing HMGB2 and SON occupancy enrichment in the 2 kbp around alternatively spliced TS sites (top) and core exons (bottom) compared to non-AS control positions (grey line). **G**, Line plots showing nucleosome occupancy deduced from Micro-C data in the 2 kbp around alternatively spliced core exons bound (bottom) or not by HMGB2 (top) in proliferating (grey) and senescent cells (green). H, Renderings of Chrom3D diploid genome models from senescent IMR90 showing TADs (beads) that carry the top 20% of CTCF (yellow), SON (light blue), or CTCF+SON signal (magenta) and AS genes from *SRRM2*-KD data. **I**, Bar plots showing clustering (±SEM) of CTCF (left), SON (middle) or CTCF+SON TADs (right) from panel H relative to randomized controls in DMSO-(grey) or ICM-treated cells (green). \**P*<0.05; unpaired two-tailed Student’s t-test. The number (*N*) of clusters deduced in each case is indicated. **J**, Bar plots showing the percent of TADs (beads) in Chrom3D models that carry most CTCF (yellow), SON (green) or CTCF+SON signal (magenta) in all different conditions. The total number of highlighted TADs (*N*) in each condition is shown. \**P*<0.05, Fisher’s exact test. **K**, Violin plots showing the exon/intron difference in GC content for differentially skipped (purple) and included exons (green) in senescence and in *SRRM2*- or *BANF1*-KD data. The number of skipping/inclusion events (*N*) and the mean *Δ*GC for ‘differential’ and ‘leveled’ exons (dotted lines; defined as in Tammer et al., 2022) are indicated. **L**, *Left*: Heat map showing the number of different AS events in genes bound by SON and HMGB2 and being differentially spliced in senescence. *Right*: GO term analysis for genes undergoing core exon (CE) skipping.

Next, we used *Whippet*^66^ to catalogue all individual AS events in our data. We began by reanalyzing data from ICM-treated IMR90, as well as from proliferating cells after *HMGB2*-KD. The former produced >5700 and the latter ~3500 AS events (using a |*Δ*Psi|>0.3 cutoff), of which 2439 were shared between the two. Just as was observed using *IsoformSwitch*, senescence induction was predominated by AS involving transcription start (TS; >35% of all AS events at |*Δ*Psi|>0.3) and end sites (TE; ~25% of all AS events at |*Δ*Psi|>0.3) (**Figure S10F** and **Table S3**). Notably, there was a strong positive correlation (ρ>0.70) between individual AS events that follow ICM treatment and those induced by *HMGB2*-KD (examples in **Figure S10G**), the main similarity of the two conditions being the formation of SICCs. For instance, when we queried genes harboring senescence-specific exon skipping, they were linked to GO terms central to the senescence program, including ‘cell cycle regulation’ and ‘chromosome organization’ (**Figure 6C**). Similarly, when we queried AS genes shared by *HMGB2*-KD and ICM, we again found key GO terms for senescence entry (**Figure S10H**).

If SICC supported the senescence splicing program, their dissolution via *SRRM2*- or *BANF1*-KD should affect AS patterns (as suggested by our *IsoformSwitch* analysis; **Figure 6A,B**). To test this, we considered all AS events with a |*Δ*Psi|>0.3. Therein, 68% of *SRRM2*- and 70% of *BANF1*-KD AS genes were also alternatively spliced upon senescence induction with ICM, the three conditions sharing 1164 AS targets (**Figure 6D**). Most strikingly, individual AS events upon *BANF1*- or *SRRM2*-KD showed a near perfect anti-correlation to the splicing events triggered by ICM (ρ<-0.70), while displaying strong positive correlation to one another (ρ>0.89; Figures 6E and **S10I**). Many of these AS events occurred at positions marked by HMGB2 and SON binding in proliferating cells (**Figure 6F**). For example, AS exons that are skipped in senescence were marked by HMGB2/SON at their 5’ intron/exon boundary in proliferating cells and lost nucleosome demarcation—a means for exon definition^67^— upon senescence entry (**Figure 6G**).

In addition to chromatin, HMGB2 was also proposed to bind RNA^68,69^, much like HMGB1 binds and regulates SASP-related mRNAs on the path to senescence^14^. We therefore used sCLIP^70^ to catalogue HMGB2 mRNA targets and crossed these with AS genes following ICM treatment (|*Δ*Psi|>0.3). This revealed an additional 287 AS events likely driven by HMGB2 loss at the RNA level, and again with a clear enrichment for alternative TS/TE events linked to GO terms central to senescence entry (**Figure S11A,B**). However, this was not also the case for CTCF mRNA targets that only included a handful of AS genes (**Figure S11C-E**). Together, these results suggest that preventing CTCF clustering broadly reverses ICM-induced AS patterns at sites marked by HMGB2 and/or SON binding on DNA and RNA targets.

Finally, we traced AS gene loci onto our Chrom3D models (**Figure 5L**) in order to gauge their spatial organization. We found that, in senescence, CTCF-rich TADs carrying AS loci clustered significantly more together and somewhat more with CTCF+SON TADs, whereas SON-rich TADs appeared to cluster more with one another than with CTCF or CTCF+SON ones (**Figure 6H,I**). At the same time, we saw significant increase in the number of CTCF-rich TADs carrying AS loci at the expense of CTCF+SON ones in senescence (**Figure 6J**). But, how does this reorganization affect AS patterns? Recent work revealed how differences in the exon-intron GC content of genes residing in the periphery compared to those residing closer to speckles correlated with different splicing outcomes. The former mostly displayed ‘differential’ architectures (i.e., larger exon-to-intron GC difference), whereas the latter mostly ‘leveled’ ones (i.e., smaller exon-to-intron GC difference^72^). As SON-bound genomic regions resided in the A1 subcompartment (**Figure S8E**) and TSA-seq–positive TADs centrally to the nucleus (**Figure 4I**), we reasoned that SICCs forming on speckles would preferentially include a specific subset of exon-intron architectures. We therefore calculated exon-intron GC content indexes for exons skipped or included upon senescence entry (|*Δ*Psi|>0.3). Skipped exons were twice as many as included ones and displayed significantly more ‘leveled’ architectures (**Figure 6K**). Notably, this bias was fully inverted upon *SRRM2*-KD and SICC dissolution, while *BANF1*-KD-dependent exons showed a similar trend (**Figure 6K**). *SRRM2*-KD-dependent exons could be linked to GO terms central to the senescence program like ‘chromosome organization in cell cycle’ or ‘cell cycle regulation’ (**Figure 6L**). This then suggests that senescence entry brings ‘differential’ exons (that are usually peripheral) closer to nuclear speckles via SICCs to induce their inclusion, while the converse applies to ‘leveled’ ones.

## DISCUSSION

The discovery of senescence-induced CTCF clusters (SICCs) in human cells reaching the endpoint of their replicative lifespan^8^ established the contribution of 3D genome reorganization in this irreversible cell fate decision^17^. At the same time, our discovery raised three important questions. How are SICCs initiated by the nuclear loss of HMGB2? What are the molecular determinants of these CTCF clusters? What functional purpose do they serve during senescence entry? Here, we addressed all three outstanding questions to show that this 3D reorganization of nuclear architecture on the path to senescence is CTCF-mediated and directs alternative splicing patterns.

Entry into senescence by normal human cells is marked by the nuclear depletion of HMGB2. HMGBs are highly abundant chromatin binders (the most abundant nuclear proteins after histones^27^) that control different legs of the senescence program^8,14^. Selective loss of HMGB2 from the nuclei of still-proliferating cells suffices to induce three major effects: a shift in favor of constitutive (HP1α-marked) heterochromatin, a global reduction in total nascent RNA output, and the formation of SICCs^8^. And, if the first effect likens the senescence-associated heterochromatin foci (SAHF) in oncogene-induced senescence^73^, the other two appear specific to replicative senescence. We can now link these two effects to explain the hitherto elusive relationship between HMGB2 loss and CTCF clustering. Despite HMGB2 and CTCF binding thousands of sites genome-wide, they display very limited overlap. Moreover, HMGB2 and CTCF do not co-purify in co-IP experiments^8^, while CTCF together with cohesin are now proposed to underlie association of chromatin with nuclear speckles in homeostasis^74^. Here, by a combination of biochemical and NGS experiments with simulations, we find that HMGB2-bound loci engage in an antagonistic relationship with CTCF-bound ones for speckle association. In the absence of HMGB2 in senescence, this balance tilts and facilitates CTCF clustering on speckles.

Is the global reduction in RNA output implicated in this? There are different aspects that need to be considered in this. First, that, upon pharmacological inhibition of transcription, nuclear speckles become larger and more round^75^ and exchange some of their components^56^. Second, that RNase A treatment of senescent nuclei does not compromise SICCs, while at the same time strengthening the interaction between CTCF and speckle components. Third, that super-resolution imaging shows SICCs forming on the surface of speckles rather than mixing inside of them. Based on our findings, we can propose that nuclear RNA acts as a general surfactant for the association of CTCF with speckles, and that global RNA reduction in senescence promotes CTCF clustering on their surface. However, it was shown that CTCF functions on chromatin, at a subset of its cognate sites, relies on (not necessarily specific) interactions with RNA^50,51^. This becomes relevant for SICC formation, as reduced RNA production in senescence would allow for more labile CTCF binding to chromatin and, thus, to a somewhat larger unbound pool. Results from Zn^+2^ manipulation experiments, as well as from simulations where we varied bound/unbound CTCF ratios (**Figure S8H**), suggest that unbound CTCF is needed for SICC emergence.

Nevertheless, SICC formation appears to nucleate on CTCF-bound chromatin (e.g., DNase I treatment of nuclear extracts abolishes CTCF-SON interaction) and speckles are now understood to associate strongly with chromatin as evidenced by biochemical experiments^76^, by the fact that SRRM2 depletion affects interactions predominantly in the A1 subcompartment^77^, and by our SON ChIPmentation profiles showing extensive chromatin linkage. Recent evidence from imaging and genomics approaches (e.g., from SPRITE^78^) also support the concept that the surface of nuclear speckles hosts a major transcriptionally active chromosomal compartment implicated in fine-tuning gene expression (reviewed in ref^79^). In fact, there exist examples, where the expression of genes is amplified upon association with speckles, including heat shock and p53-responsive genes^35,80,81^. It is therefore not unreasonable to propose that gene loci kept in close proximity to nuclear speckles via SICCs might benefit from their sustained positioning within a highly reactive niche in the otherwise transcriptionally subdued senescent nucleus.

Nuclear speckles are now shown to be multi-layered, non-randomly organized condensates^29,45,82,83^. At the same time, it was recently shown that the N-terminal and DNA-binding domains of CTCF can also form condensates *in vitro* and inside of cells, which may facilitate its insulatory role^39^. We can now complement these findings by showing that CTCF clustering is sensitive to 1,6-hexanediol and can be competed out by specific, albeit short intrinsically disordered domains, like the acidic tail of HMGB2. Therefore, SICCs dissplay condensate-like behavior while forming on the surface of speckles with BANF1 and SRRM2 being necessary for clustering. We propose that both of these components are repurposed by senescent cells. BANF1 is required for accurate segregation of mitotic chromosomes^44^, but in the context of irreversible cell cycle arrest a subset of BANF1 molecules is moved from the periphery to the nuclear interior to act as a ‘molecular glue’ in SICCs. SRRM2 is a *bona fide* splicing factor, the RNA-binding domain of which suffices for CTCF clustering. However, senescent cells need to execute a radically different gene expression program and do this in the context of reduced overall transcriptional potency. In homeostasis, nuclear speckles can induce splicing and accelerate the processing of RNAs located in their vicinity, while also buffering the availability of its components between the different nuclear compartments^33,84^. Here, we show that disruption of SICCs via SRRM2 or BANF1 depletion leads to the near-complete reversal of splicing patterns induced upon senescence entry. This is likely due to the repositioning of SICC-associated loci away enough from speckles. Interestingly, we find that exons skipped the most upon senescence entry are ‘leveled’ ones that preferentially reside close to speckles in homeostasis, and those included the most are ‘differential’ ones that tend to be peripheral^72^. These effects imply that SICCs will tether in the vicinity of speckles loci that are not usually found there in proliferating cells, thereby changing their splicing fates—and, thus, SICC dissolution undoes these splicing choices.

In summary, we provide evidence linking both large- and fine-scale 3D genome reorganization with the implementation of a specific splicing program underlying cellular aging. This does not only represent a striking paradigm of the interplay between overall nuclear architecture and gene expression regulation beyond transcription, but may also provide an additional entryway into senescence modulation. As the load of senescent cells in ageing tissues and organs is causally implicated in health- and lifespan extension^2^, interfering with nuclear speckle and SICC physiology might provide new means for delaying senescence or for alleviating part of its detrimental effects during the course of ageing.

## ACKNOWLEDGEMENTS

We thank Vassilis Roukos for critical reading of the manuscript. This study was supported by the German Research Foundation (DFG) via the Priority Programs SPP2202 (project 422389065 awarded to A.P. and 507778679 awarded to A.M.O.) and SPP2191 (project 506296585 awarded to A.P.), the Collaborative Research Center 1565 (projects 469281184 awarded to A.P. and 469281184 awarded to A.M.O.), the TRR81/3 (project 109546710 awarded to A.P.), and by the Lower Saxony Ministry for Science and Culture (MWK) via the SPRUNG (project 76211-1267/2023 awarded to A.P.) and BEREIT grants (project 2019-00298 awarded to A.P.). L.K. is supported by an ERC Consolidator Grant (TRANSCEND), and A.M.O. is supported by an ERC Starting Grant (3D-REG). S.P., V.V.-M., and S.R. are supported by the IMPRS Genome Science PhD program, and Y.Z. by the IMPRS-MolBio MSc/PhD program of the University of Göttingen.

## AUTHOR CONTRIBUTIONS

S.P., I.L., A.M., I.P., and K.S. performed experiments. V.V.-M. performed NGS data analysis and generated Chrom3D models. Y.Z. performed splicing analyses. S.R. and A.M.O. contributed ChIPmentation data. A.S. and C.N. analysed Micro-C. D.B., Y.K., and L.K. produced CLIP data. M.B. performed MD simulations. A.P. designed the study and secured funding. S.P. and A.P. compiled the manuscript with input from all co-authors.

## DATA AVAILABILITY

All NGS data generated in this study is available via the NCBI Gene Expression Omnibus (GEO) under accession numbers GSE269856 [RNA-seq], GSE269857 [eCLIP], GSE269858 [ChIPmentation]. HMGB2 sCLIP data was retrieved from GSE171782^14^, and Micro-C and CTCF CUT&Tag data from GSE238256^38^.

## METHODS

### Cell Culture

Proliferating primary lung fibroblasts (IMR90) isolates (I79 and I83, passage 5; Coriell) were grown in MEM (M4655, Sigma-Aldrich) supplemented with 1x non-essential amino acids and 10% FBS under 5% CO_2_. Senescence was induced after treating the cells for up to 6 days with 10μM inflachromene (ICM; Cayman Chemicals) as described^38^. Where indicated, cells were treated with 6% 1,6-hexanediol (Sigma-Aldrich) for 1 min at room temperature.

### Protein co-immunoprecipitation

Approximately 6×10^6^ proliferating or ICM-treated IMR90 were gently scraped and pelleted for 5 min at 700x *g*, resuspended in 500 μl ice-cold lysis buffer (20 mM Tris-HCl pH 8.0, 1% NP-40, 150 mM NaCl, 2 mM EDTA pH 8.0) supplemented with 1x protease inhibitor cocktail (Roche), and incubated for 30 min on ice, followed by three cycles of sonication (30 sec *on*, 30 sec *off*; low input on a Bioruptor Pico) and benzonase treatment, before centrifugation for 15 min at > 20,000x *g* to pellet cell debris and collect the supernatant. During these steps, 30 μl protein-G magnetic beads (Active Motif) and 10 μg CTCF antibody (Active motif, 61311) were incubated for 2 h at 4°C on a rotor, before the beads were captured on a magnetic rack (Active Motif) and added to the lysates for incubation at 4°C overnight under rotation. The next day, the beads were washed 4x with 900 μl ice-cold wash buffer I (50 mM Tris, 0,05% NP-40, 50 mM NaCl), 2x with 500 μl ice-cold wash buffer II (150 mM NaCl, 50 mM Tris), recaptured on the magnetic rack and the supernatant discarded. Proteins remaining associated with the beads were pre-digested in 50 μl elution buffer (2 M urea, 50 mM Tris pH 7.5, 1 mM DTT, 50 ng trypsin) for 30 min at room temperature with gentle agitation. Following addition of 50 μl digestion buffer (2 M Urea dissolved in 50 mM Tris pH 7.5, 5 mM chloroacetamide) and incubation for 30 min at room temperature, another 50 μl of elution buffer supplemented with 50 ng of LysC and 100 ng of trypsin were added to each tube and allowed to be digested overnight at room temperature. Next day, the digestion was stopped by adding 1 μl trifluoroacetic acid, peptides of each experiment were split in half, purified on two C18 stage tips, and all replicates were analyzed on a Q-Exactive platform (Thermo Fisher; full results in **Table S1**). For the data in **Figure S6E**, benzonase treatment was replaced by treatment with RNase A or DNase I (100 µg/ml) and, following the same washes as above, beads were resuspended in 20 µl 1X Laemmli buffer and boiled at 95°C for 10 min to elute proteins before western blot analysis.

### Immunofluorescence, image acquisition and quantification

IMR90 grown on glass acid-etched coverslips were fixed in 4% PFA/PBS for 10 min at room temperature. After washing once in PBS, cells were permeabilized with 0.5% Triton-X/PBS for 5 min at room temperature, blocked with 1% BSA for 1 h, and incubated with a 1:500 dilution of the appropriate primary antibody for 1 h at room temperature (anti-CTCF, Active Motif 61331; anti-SRRM2, Thermo Fisher PA5-59559; anti-SC35, Novus Biologicals NB100-1774; anti-HMGB1, Abcam ab190377-1F3). The primary antibody was washed 2x with PBS for 5 min per wash and cells were incubated with secondary antibodies (diluted in 0.5% BSA/PBS) at room temperature in the dark for 1 h at 1:1000 dilution (anti-rabbit Alexa488, Abcam ab150077; anti-mouse Cy3, Abcam ab97035). Cells were then washed with 2x with PBS for 5 min per wash before ProLong^TM^ Gold antifade reagent with DAPI (#P36931) was added to them. All widefield images were acquired on a Leica DMI8 microscope with an HCX PL APO 63x/1.40 (Oil) objective via the LASX software. Super-resolution images were acquired on an Abberior STEDYCON microscope with a 100x Plan SuperApochromat 1.4 Oil objective. Quantification of nuclear fluorescence were performed by drawing a mask based on DAPI staining and calculating mean intensity per area under this mask. Co-localization was assessed using a FIJI plugin (https://imagej.nih.gov/ij/plugins/rgb-profiler.html).

### RNA isolation and RT-qPCR analysis

IMR90 were washed with 1x PBS and harvested directly in Trizol (Invitrogen, 15596018). RNA was isolated from this lysate by using the Direct-zol RNA MiniPrep Kit (Zymo Research, R2052:1074069) including an on-column DNase I treatment step according to the manufacturer’s instructions. Then, ∼1 μg total RNA was used for synthesizing cDNA using the SuperScript First-Strand Synthesis System for RT-PCR (Invitrogen, 11904018). For the qPCR, the qPCRBIO SyGreen Mix (PCR Biosystems, PB20.14-51) was used on 2 ng of cDNA with reactions mixed in 384-well skirted PCR plates (FrameStar, 4ti-0385) and loaded onto a qTOWER^3^ Real time thermocycler (Analytik Jena). All qPCR primers are listed in **Table S4**.

### siRNA-mediated knockdown

IMR90 were seeded at ∼35,000 cells/cm^2^ the day of transfection and RNAiMAX Transfection Reagent (Invitrogen, 13778075) was used for preparing siRNA mixtures and delivering to the cells according to the manufacturer’s instructions. Knockdown efficiency was assessed 48h after transfection using RT-qPCR and immunostainings, and RNA isolated from cells was used for cDNA library construction using the TruSeq kit (Illumina) before paired-end sequencing on a NovaSeq6000 platform (Illumina). Details for all siRNAs are provided in **Table S5**.

### Overexpression experiments

Doxycycline-inducible overexpression in IMR90 was achieved via piggyBac transposition as described previously^8^. The SRRM2 RNA binding and C-terminal IDR domains, as well as the full length and all HMGB2 subparts were amplified and cloned from cDNA. Following validation by Sanger sequencing, they were separately subcloned into the DOX-inducible KA0717 expression vector to generate Venus fusions. Each construct was co-transfected into IMR90 together with transactivator and transposase-encoding vectors (KA0637 and SBI Biosciences #PB200PA-1, respectively) at a DNA mass ratio of 10:1:3 using Lipofectamine® LTX reagents (InvitrogenTM, #56532) as per manufacturer’s instructions. Stable, transgene-positive, proliferating IMR90 were selected using 350 μg/ml G418 (Sigma Aldrich), expression was induced using doxycycline for 24h, and IMR90 carrying the empty Venus vector served as a control.

### ChIPmentation experiments

ChIPmentation for SON and HMGB2 was performed in three biological replicates per condition. We optimized a published protocol^85^ to improve the signal-to-noise ratios in the data. In brief, aliquots of 1 million IMR90 were crosslinked by double fixation with 1.5 μM EGS for 20 min and 1% formaldehyde (Electron Microscopy Sciences) for 10 min at room temperature, quenched with ice-cold glycine (Sigma-Aldrich, G7126) at a final concentration of 125 mM, and washed 2 times with ice-cold PBS. Fixed cells were first gently lysed with Farnham lysis buffer (5 mM PIPES pH 8.0, 85 mM KCl, 0.5% NP-40) to isolate nuclei, followed by nuclear lysis with 0.5% SDS buffer (10 mM Tris-HCl pH 8, 1 mM EDTA, 0.5% SDS). Then, chromatin was fragmented, using optimized conditions involving a titrated amount of MNase (2.5 Kunitz units) and gentle sonication (time: 1 min, duty factor: 2 %, peak incident power: 105 W, cycles per burst: 200). For immunoprecipitation, a mixture of Protein A (10008D, Invitrogen) and Protein G Dynabeads (10003D, Invitrogen) in 1:1 ratio was blocked with 0.5 % BSA, washed and pre-incubated with 2 µg of the relevant primary antibody for 6 h at 4 °C. Sheared chromatin was diluted in IP buffer (10 mM Tris-HCl pH 8, 1 mM EDTA, 150 mM NaCl, 1% Triton-X-100, 1X Protein Inhibitor Cocktail), and added to the bead-bound antibodies. Samples were incubated overnight on a rotator at 4 °C, and washed with stringent buffers on the next day. Sequencing adapters were added to the bead-bound DNA by tagmentation with the Tagment DNA Enzyme and Buffer Kit (Illumina). After washes, bead-bound samples were de-crosslinked overnight at 65 °C in the presence of proteinase K (0.2 mg/mL) and cleaned with 1.8x volume of Mag-Bind TotalPure beads (Omega Bio-Tek). Tagmented DNA was used for library preparation with the NEBNext High-Fidelity 2X PCR Master Mix (NEB). The quality of the libraries was assessed with a fragment analyzer and the libraries were sequenced on the Illumina sequencing platform with 75 cycles paired-end reads. Peak lists are included in **Table S6**.

### CLIP and data analysis

Enhanced cross-linking with immunoprecipitation (eCLIP) was performed as previously described^54^. In brief, ∼20×10^6^ IMR90 were UV-crosslinked (400 mJ/cm^2^ constant energy), lysed in iCLIP lysis buffer, and sonicated (on a BioRuptor Pico). Lysates were treated with RNase I (Ambion, AM2294) to fragment RNA, after which CTCF protein-RNA complexes were immunoprecipitated using the relevant antibody (anti-CTCF, Active Motif 61331). In parallel to the IP, an “input” library was generated for each replicate for which no antibody was used. Stringent washes were next performed, during which RNA was dephosphorylated with FastAP enzyme (Fermentas) and T4 PNK (NEB, M0201S). Subsequently, a 3’ RNA adaptor oligonucleotide was ligated onto the RNA using a T4 RNA ligase (NEB, M0242S). Protein-RNA complexes were separated on an SDS-polyacrylamide gel electrophoresis gel and transferred to nitrocellulose membranes, and RNA was isolated off the membrane. After precipitation, RNA was reverse-transcribed with AffinityScript reverse transcriptase (Agilent, 600107), free oligos were removed using ExoSap-IT (Thermo Fisher Scientific, 78201.1.ML), and a 3’ DNA adaptor was ligated onto the cDNA product. Libraries were then amplified with 2x Q5 PCR mix (NEB), paired-end sequenced on a HiSeq3000 platform (Illumina) and analyzed via the eCLIP pipeline (https://github.com/YeoLab/eCLIP). All HMGB2 and CTCF CLIP targets are listed in **Table S7**.

### Splicing analysis

Paired-end RNA-seq reads were mapped to human reference genome build hg38 using STAR ver. 2.7.3a^86^ with default parameters, uniquely mapped gene counts quantified using HTSeq ver. 0.12.4^87^, and normalizes using the RUVs module of the RUVseq package ver. 1.30.0^88^. Differentially expressed genes were defined via the DEseq2 ver. 1.36.0 package using its default Wald test^89^ and deemed as DEGs when *P*_adj_ <0.05 and absolute log_2_FC >0.6. Alternatively spliced (AS) transcripts and AS events were quantified using Whippet ver. 1.6^66^ and catalogues once they had a probability >0.9 and an absolute ΔPsi >0.1. The GC content difference between exons and their flanking introns was based on the method described previously^72^ using bedtools ver. 2.29.1^90^, whereby the mean intronic GC content is subtracted from the exonic GC content value. For this, only the 15 nt proximal to the exon were considered in these calculation for introns >150 nt (otherwise the whole intronic length was used), and splice sites (defined as the first 2 and the last 3 nt of each exon, and the last 20 nt of its upstream and the first 6 nt of its downstream intron) were excluded. The 25,000 exons with the highest and the lowest GC content difference genome-wide were extracted from all the expressed exons that are longer than 75 nt to calculate the reference “differential” and “leveled” GC content difference for IMR90. Finally, for genome-wide mRNA isoform analysis, transcripts were quantified using salmon ver. 1.5.0^91^, and functional isoform switches annotated using isoformSwitchAnalyzeR ver. 2.2.0 (with an isoform fraction change of >0.3)^65^. Functional GO term enrichment analysis of genes was performed via Metascape^92^.

### Micro-C data analysis and modeling via Chrom3D

Micro-C data from proliferating and ICM-treated IMR90^38^ were analysed using the Dovetail Genomics pipeline (https://micro-c.readthedocs.io/en/latest/fastq_to_bam.html) and used an input for Chrom3D modeling. In brief, read pairs were mapped to the human reference genome hg38 using BWA, after which low mapping quality (<40) reads and PCR duplicates were filtered out using the *MarkDuplicates* function in Picard tools (v2.20.7), and read coverage tracks (BigWig) were generated and normalized with the RPCG parameter using the *bamCoverage* function of deepTools2 v3.5.1^93^. For Chrom3D simulations^61^, TADs were called at 10-kbp resolution using HiCPro v2.11.4^94^, and intra-TAD interactions specified according to Micro-C signal. Association with LADs was considered as described in the Chrom3D manual for the whole diploid genome (https://github.com/Chrom3D) and assigned to each chromosome; constitutive LADs were inferred from the LAD Atlas (https://osf.io/dk8pm/wiki/home/). In the end, a .*gtrack* file (Chrom3D input) for chromosome visualization was produced using existing scripts (https://github.com/Chrom3D/preprocess_scripts). Next, a .*BED* file specifying the genomic positions of the TADs (1 TAD = 1 bead) was created, and any gaps between them were filled by non-TAD beads of the appropriate size. Finally, .*gtrack* files corresponding to each cluster were merged and inputted in Chrom3D, using 1,000,000 iterations (-n), a nuclear radius of 5 (-r), and a scale total volume of the beads relative to the volume of the nucleus set to 0.15 (-y). For whole genome visualizations that take into account interchromosomal interactions, LADs, TADs, and Micro-C matrices were used for the production of a diploid .*gtrack* file using default parameters; chromosomes Y and M were removed. IDs of beads containing the top 20% of CTCF and/or SON signal from proliferating or ICM-treated cells were identified and colored using the *processing_scripts/color_beads.py* script and the *blend* flag to maintain coloring. All 3D models were visualized in Chimera-X v1.3^95^. For the clustering analysis, we employed a DBSCAN-inspired density-based clustering non-parametric algorithm to identify clusters of TADs in the Chrom3D models. To tackle the more complex task of clustering the monomers of a polymer with Euclidean distance-based criteria, in addition to the classical DBSCAN algorithm, we excluded the possibility for two TADs to be in the same cluster only because they are *n* TADs away from each other along the chromosome, with *n* being an input parameter to set. For the presented analysis *n* has been set to 2. The choice of the distance cutoff was driven by finding the trade-off between identifying enough clusters but with a large enough number of members each, to have enough statistics power. From cluster statistics analysis, we selected a cutoff distance of 0.3 as the good compromise. Also, the minimum number of members to call a cluster was set to 5. In the Chrom3D models, TADs are classified as ‘CTCF’, ‘SON’, ‘CTCF+SON’ or ‘other’ depending on signal enrichment. Once clusters were identified, we calculated their TAD-type composition and compared with the expected composition had they just been randomly extracted from the entire pool of TADs. The (log_2_) fold change of these two quantities gives a quantitative indication of how many clusters are enriched for a given TAD type with respect to random expectation. For significance estimation we employed the Mann-Whitney U-rank test.

### Molecular Dynamics simulations

Molecular dynamics simulations were performed via the multipurpose EspressoMD package^96^ as described^62^. Briefly, individual proteins are represented by beads interacting via phenomenological force fields and the chromatin fiber as a chain of beads connected by bonds. The position of every bead in the system evolved according to the classical Langevin differential equation encoding Newton’s laws in the case of thermal bath with friction *γ* due to an implied solvent, and in the presence of forces between beads given by energy potential functions. By dynamically forming and dissolving protein-chromatin bonds, this framework simulates RNAPII traversing of genes during transcription and chromatin loops extruding by the cohesin complex and CTCF protein for the purpose of 3D chromatin folding. For the present study, we added to our previous framework the two key factors HMGB2 and SRRM2, modeled as two new binder classes. Along with CTCF, all three proteins were given the ability to phase separate due to their prominent intrinsically disordered regions via a Lennard-Jones potential (LJ-contacts) encoding the multivalent character of such interactions. In addition, driven by our coIP-MS results, CTCF and HMGB2 proteins were both allowed to form multiple LJ-contacts with SRRM2 proteins, but have no interaction with each other. This experiment-driven set up provides the force background allowing for CTCF to form clusters when HMGB2 is depleted, since the former in this condition is more likely to colocalize with SRRM2 clusters. In fact, SRRM2 molecules are given higher phase separation affinity due to their vast IDRs, hence always form clusters representing nuclear speckles. For our model, we selected a 3-Mbp region (chr9:112-115 Mbp) mostly devoid of lamina-associated domains (LADs) and rich in CTCF, HMGB2, and enhancer marks. As before, H3K27ac CUT&Tag peaks were encoded as sites for RNAPII binding; similarly, CTCF CUT&Tag and HMGB2 ChIPmentation peaks were sites binding CTCF and HMGB2 beads, respectively. The harmonic potential bond between binders and chromatin binding sites has an exclusive nature, meaning that, once formed, the binding site involved cannot form another bond with another binder of the same class, while the binder involved cannot form another bond with another chromatin binding site. Four genes are encoded in the region and are subjected to active transcription by RNAPII. The presence of chromatin-bound HMGB2 allows RNAPs to bind a nearby promoter with 30% higher probability, thus modeling HMGB2 function and producing an ∼30% reduction of transcription in the modeled region upon its depletion as previously seen experimentally^8^.

### Statistical testing

*P*-values associated with Student’s *t*-tests and Fischer’s exact tests were calculated using GraphPad (https://graphpad.com/) while those associate with Wilcoxon-Mann-Whitney tests were calculated in R. Unless otherwise stated, *P-*values <0.05 were deemed as statistically significant.

## SUPPLEMENTARY INFORMATION

**Figure S1.**
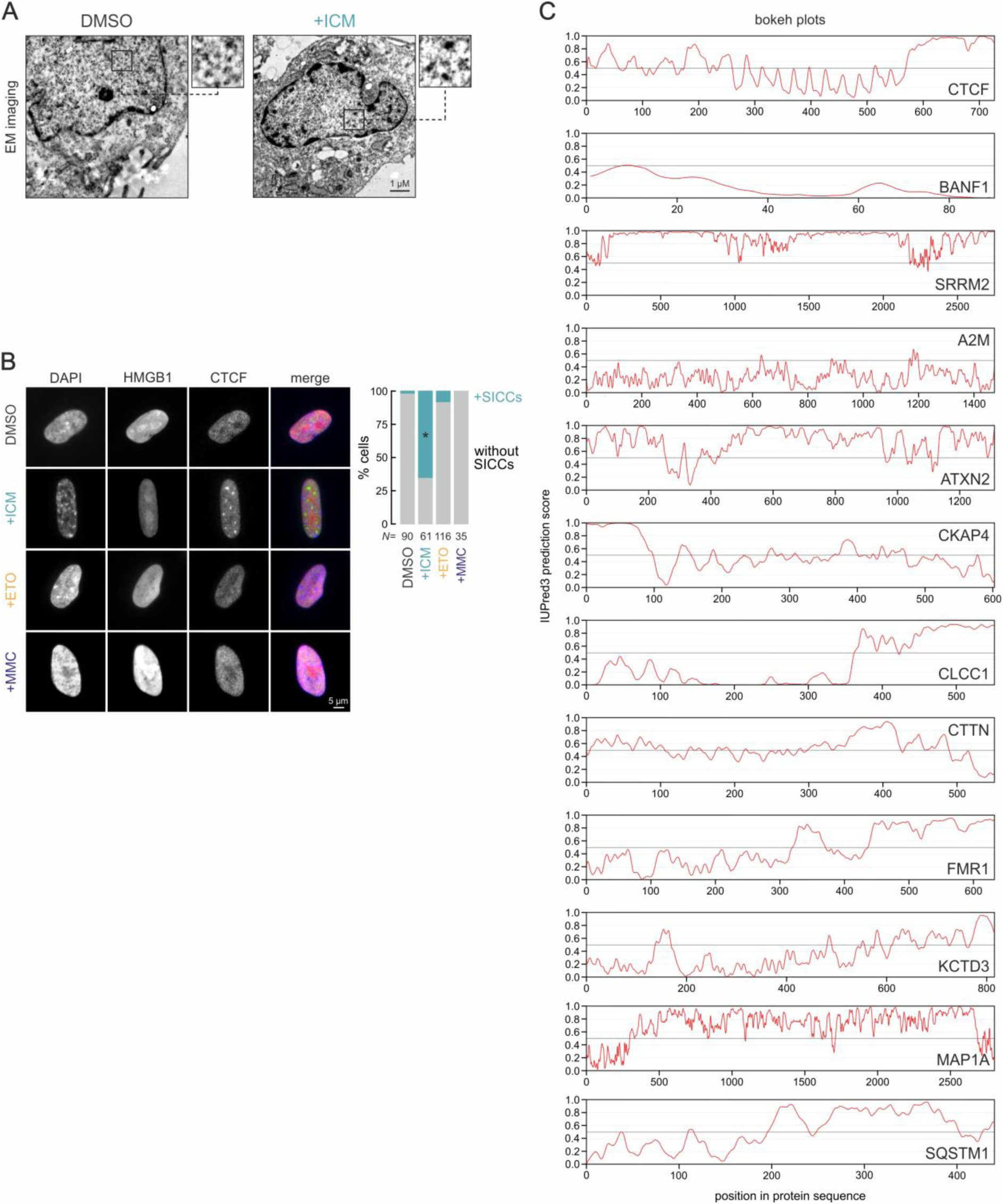
SICC formation and CTCF protein interactors in senescence (*linked to* Figure 1). **A**, Representative electron microscopy images from proliferating (DMSO) and senescent IMR90 (+ICM) immunostained for CTCF. Magnified regions (rectangles) show CTCF clustering in senescence. **B**, *Left*: Representative images of proliferating (DMSO), senescent (+ICM), etoposide-(+ETO) and mitomycin-treated IMR90 (+MMC) immunostained for HMGB1 and CTCF and counterstained by DAPI. *Right*: Bar plots of the percent of cells displaying SICCs; *N* is the number of cells quantified per condition. \**P*<0.05, Fisher’s exact test. **C**, Line plot showing the predicted IUPred3 disorder score along CTCF and 11 of its interactors from co-IP-mass spectrometry data (see **Table S1**).

**Figure S2.**
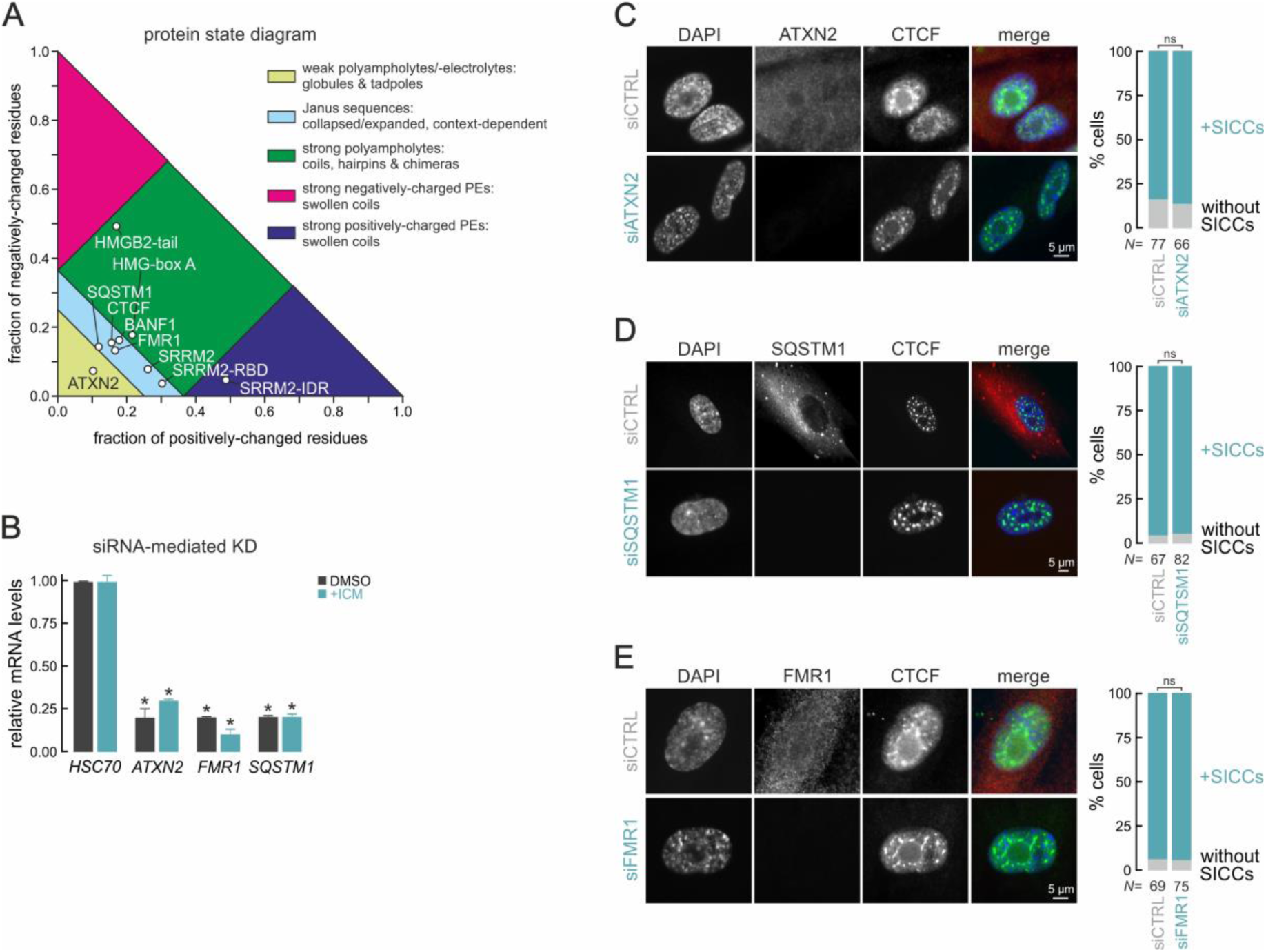
Functional testing of SICC dependence on senescence-specific CTCF interactors (linked to Figure 1). **A**, Protein state diagram summarizing the physicochemical profile of CTCF, its top tested interactors (ATXN2, BANF1, FMR1, SQSTM1, SRRM2, SRRM2-RBD, and SRRM2 C-terminal IDR), as well as of full length HMGB2 and its subcloned parts (HMGB2C-terminal acidic tail, HMG-box A). **B**, Bar plots showing changes in mRNA levels upon siRNA-mediated *ATXN2*-,-*FMR1*- or *SQSTM1*-KD in proliferating (DMSO) and senescent IMR90 (+ICM) relative to *HSC70* controls. \**P*<0.01, unpaired two-tailed Student’s t-test. **C**, *Left*: Representative images of senescent IMR90 immunostained for CTCF and ATXN2, and counterstained by DAPI at 48 h after transfection with non-targeting (siCTRL) or *ATXN2* siRNAS (siATXN2). *Right*: Bar plots showing the percentage of cells displaying SICCs; *N* is the number of cells quantified per condition. ns: *P*>0.05, Fisher’s exact test. **D**, As in panel C, but using siRNAs targeting *SQSTM1*. E, As in panel C, but using siRNAs targeting *FMR1*

**Figure S3.**
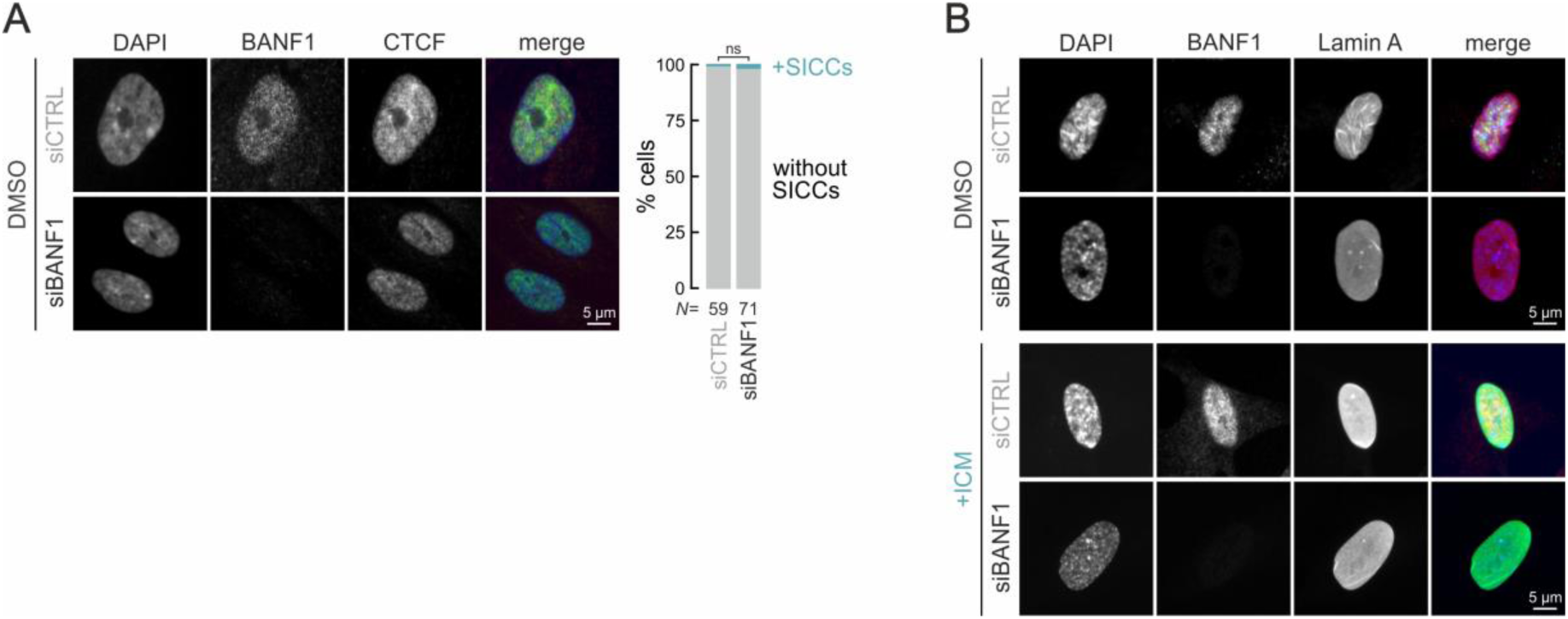
Effects of BANF1 depletion from IMR90 cells (linked to Figure 2). **A**, *Left*: Representative immunofluorescence images of proliferating IMR90 (DMSO) stained for CTCF and BANF1, and counterstained by DAPI at 48 h after transfection with non-targeting (siCTRL) or *BANF1* siRNAS (siBANF1). *Right*: Bar plots showing the percentage of cells displaying SICCs; *N* is the number of cells quantified in each condition. ns: *P*>0.05, Fisher’s exact test. **B**, *Left*: Representative immunofluorescence images of proliferating (DMSO) and senescent IMR90 (+ICM) stained for BANF1 and Lamin A, and counterstained by DAPI at 48 h after transfection with non-targeting (siCTRL) or *BANF1* siRNAS (siBANF1).

**Figure S4.**
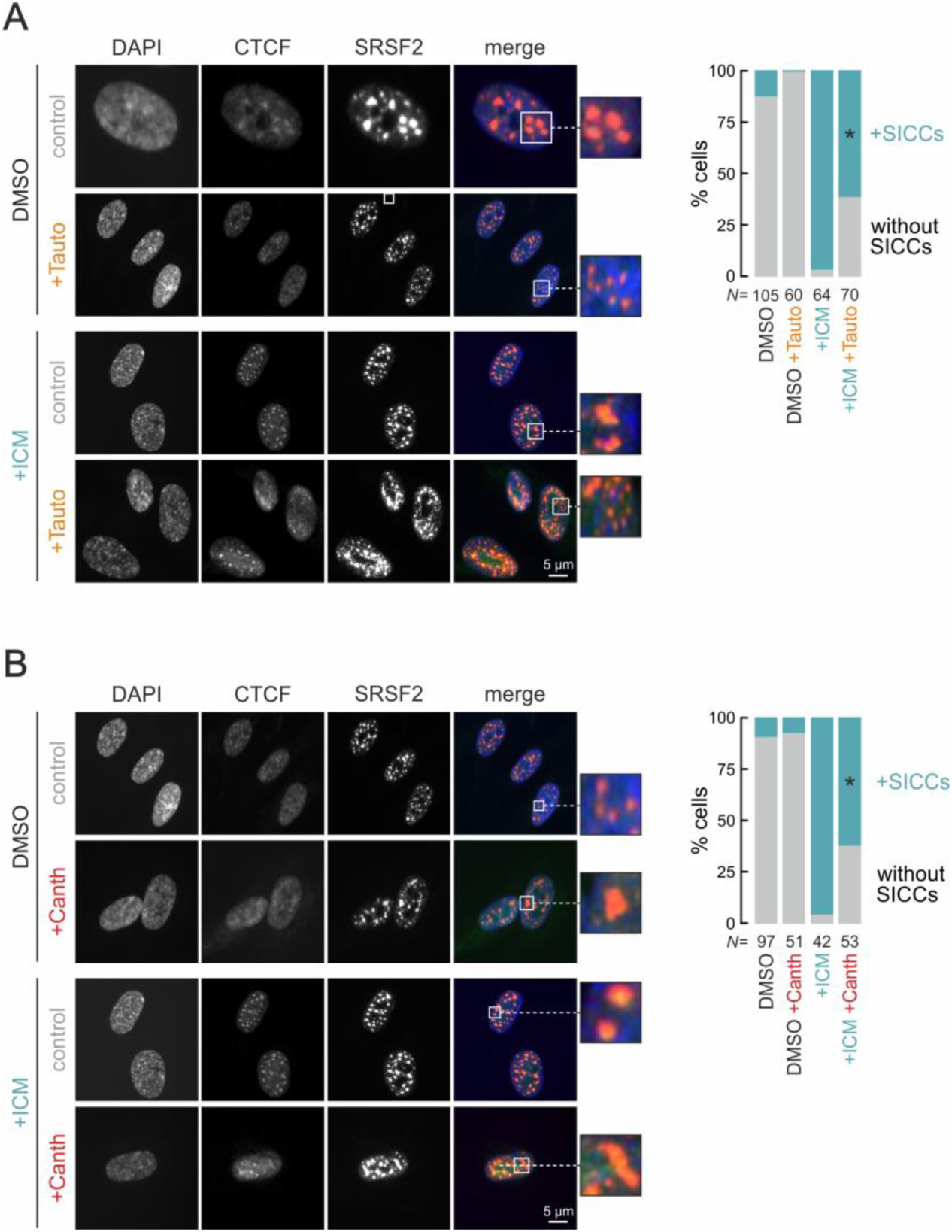
SICCs follow nuclear speckle dispersal and fusion (*linked to* Figure 3). **A**, *Left*: Representative immunofluorescence images of proliferating (DMSO) and senescent IMR90 (+ICM) stained for CTCF and SRSF2, and counterstained by DAPI following treatment or not with tautomycetine to induce speckle dispersion. *Right*: Bar plots showing the percentage of cells displaying SICCs; *N* is the number of cells quantified in each condition. \**P*<0.05, Fisher’s exact test. **B**, As in panel A, but using cantharidin to induce speckle fusion.

**Figure S5.**
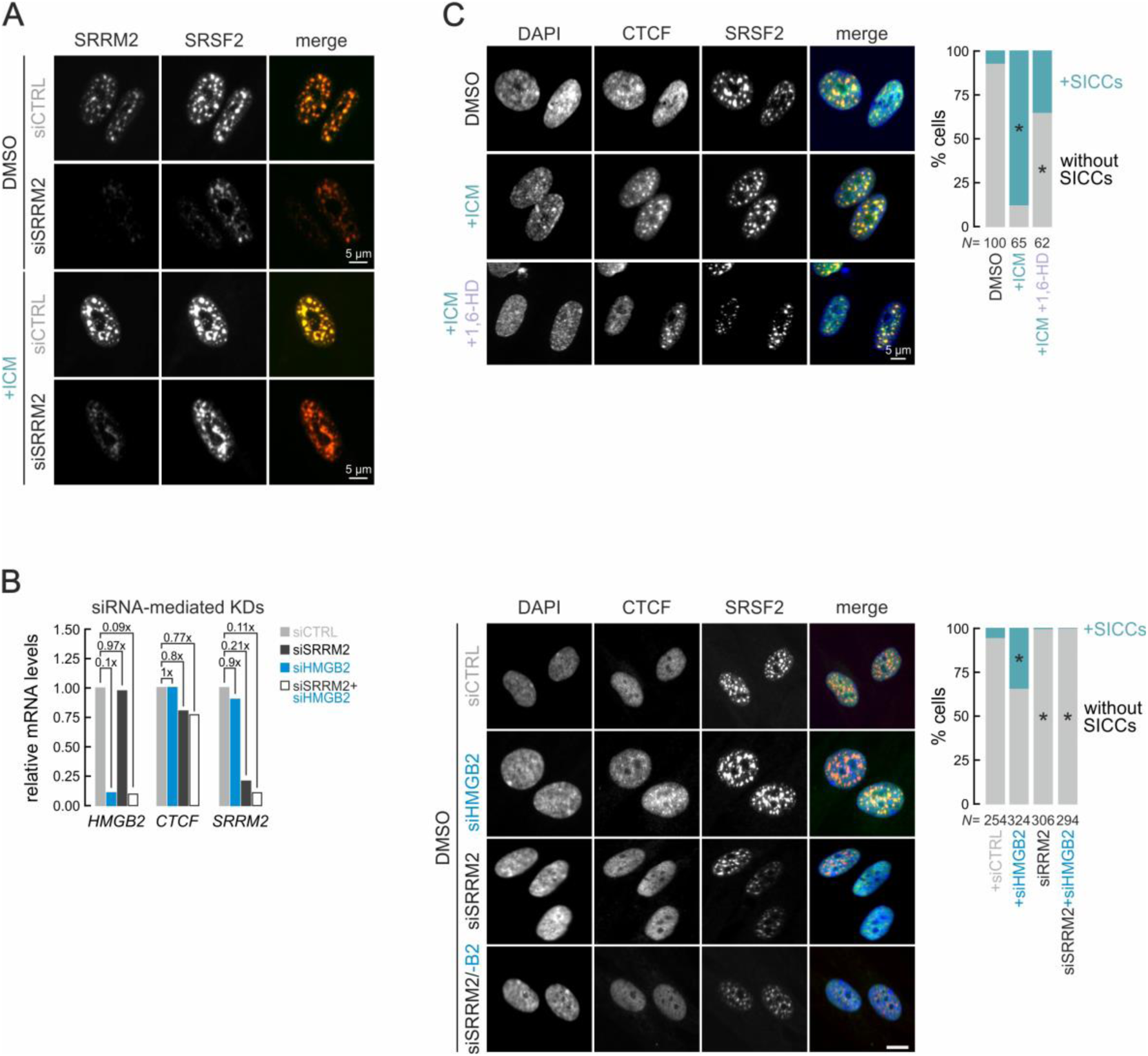
SRRM2 depletion and 1,6-hexanediol treatment does not dissolve nuclear speckles (*linked to* Figure 3). **A**, Representative immunofluorescence images of proliferating (DMSO) and senescent IMR90 (+ICM) stained for SRRM2 and SRSF2, and counterstained by DAPI 48 h after transfection with non-targeting (siCTRL) or *SRRM2* siRNAs (siSRRM2). **B**, *Left*: Bar plots showing changes in mRNA levels upon siRNA-mediated *HMGB2*- and/or *SRRM2*-KD in proliferating IMR90 (DMSO) relative to *HSC70* controls. * *Middle*: Representative immunofluorescence images of proliferating IMR90 (DMSO) stained for CTCF and SRSF2, and counterstained by DAPI at 48 h after transfection with non-targeting (siCTRL), HMGB2-(siHMGB2), *SRRM2*-(siSRRM2) or *HMGB2-* plus *SRRM2*-targeting siRNAs (siSRRM2/-B2). *Right*: Bar plots showing the percent of cells displaying SICCs. *N*, the number of cells quantified in each condition; \**P*<0.05, Fisher’s exact test. **C**, As in panel B, but for proliferating (DMSO), senescent (+ICM) and senescent IMR90 treated with 6% 1,6-hexanediol for 1 min (+ICM +1,6-HD). *N* is the number of cells quantified in each condition; \**P*<0.05, Fisher’s exact test.

**Figure S6.**
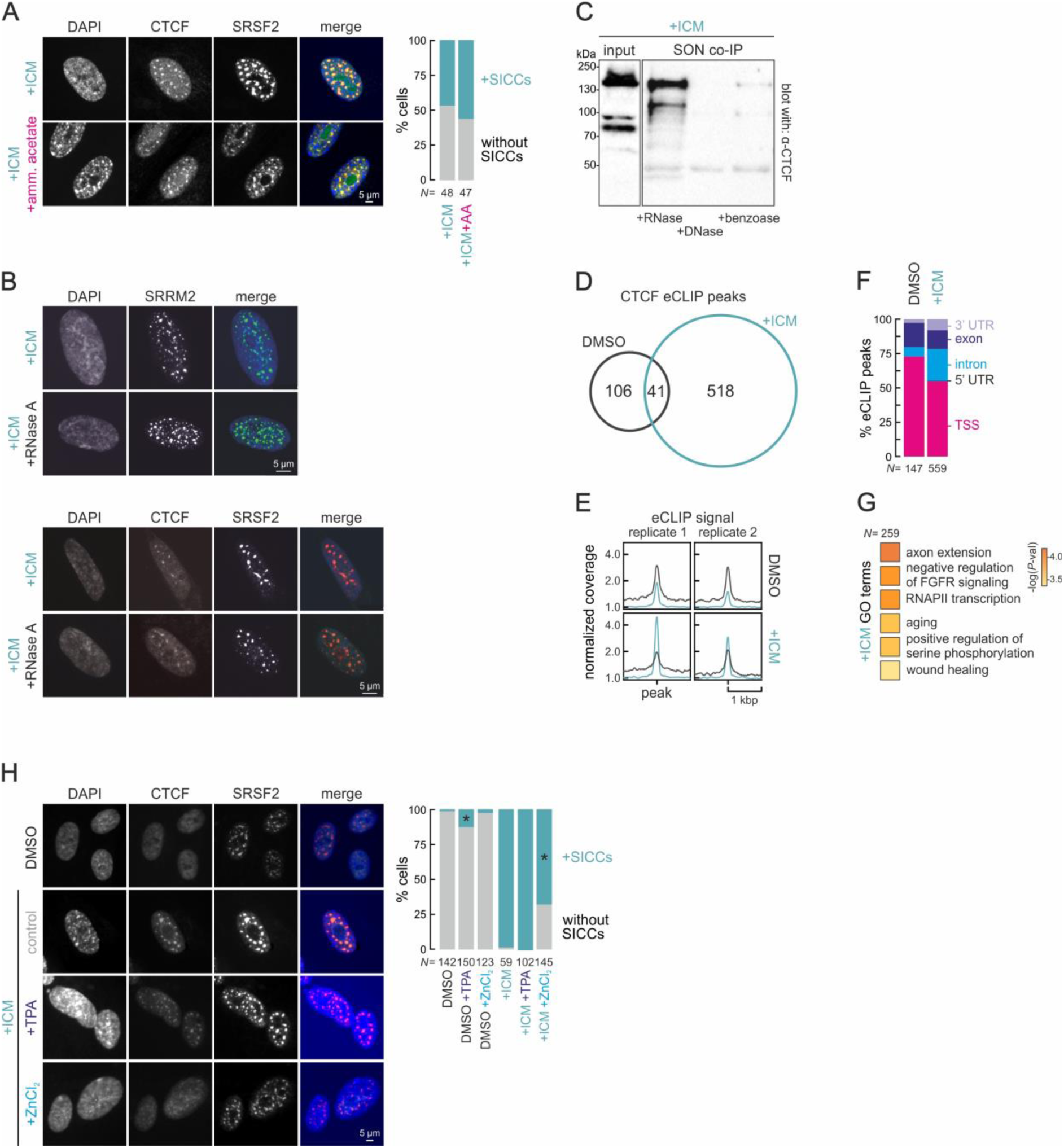
SICCs are resistant to ammonium acetate, RNase A, and zinc chelation (*linked to* Figure 3). **A**, *Left*: Representative immunofluorescence images of senescent IMR90 (+ICM) treated with 100 mM ammonium acetate for 10 min, stained for CTCF and SRSF2, and counterstained by DAPI. *Right*: Bar plots showing the percent of cells with SICCs. *N*, the number of cells quantified; \**P*<0.05, Fisher’s exact test. **B**, As in panel A, but for senescent IMR90 (+ICM) treated or not with RNase A for 20 min and stained for SRRM2 (top) and for CTCF and SRSF2 (bottom), and counterstained by DAPI. **C**, Western blot analysis of SON co-IP experiments performed using benzonase, (RNase-free) DNase I or RNase A to treat lysates before blotting with an antibody against CTCF. The 130-KDa CTCF input band provides a control. **D**, Venn diagram of the overlap of CTCF eCLIP peaks from proliferating (DMSO) and senescent IMR90 (+ICM). **E**, Mean eCLIP signal from two replicates in the 2 kbp around proliferating (DMSO) and senescent peaks (+ICM). **F**, Bar plots showing the distribution of CTCF eCLIP peaks in proliferating (DMSO) and senescent IMR90 (+ICM). G, Heat map showing GO term enrichment for the 259 CTCF-bound mRNAs in senescence. **H**, As in panel A, but for proliferating (DMSO) and senescent IMR90 (+ICM) treated with 50 µM of a zinc chelating agent (TPA) or with 30 µM ZnCl_2_ for 30 min.

**Figure S7.**
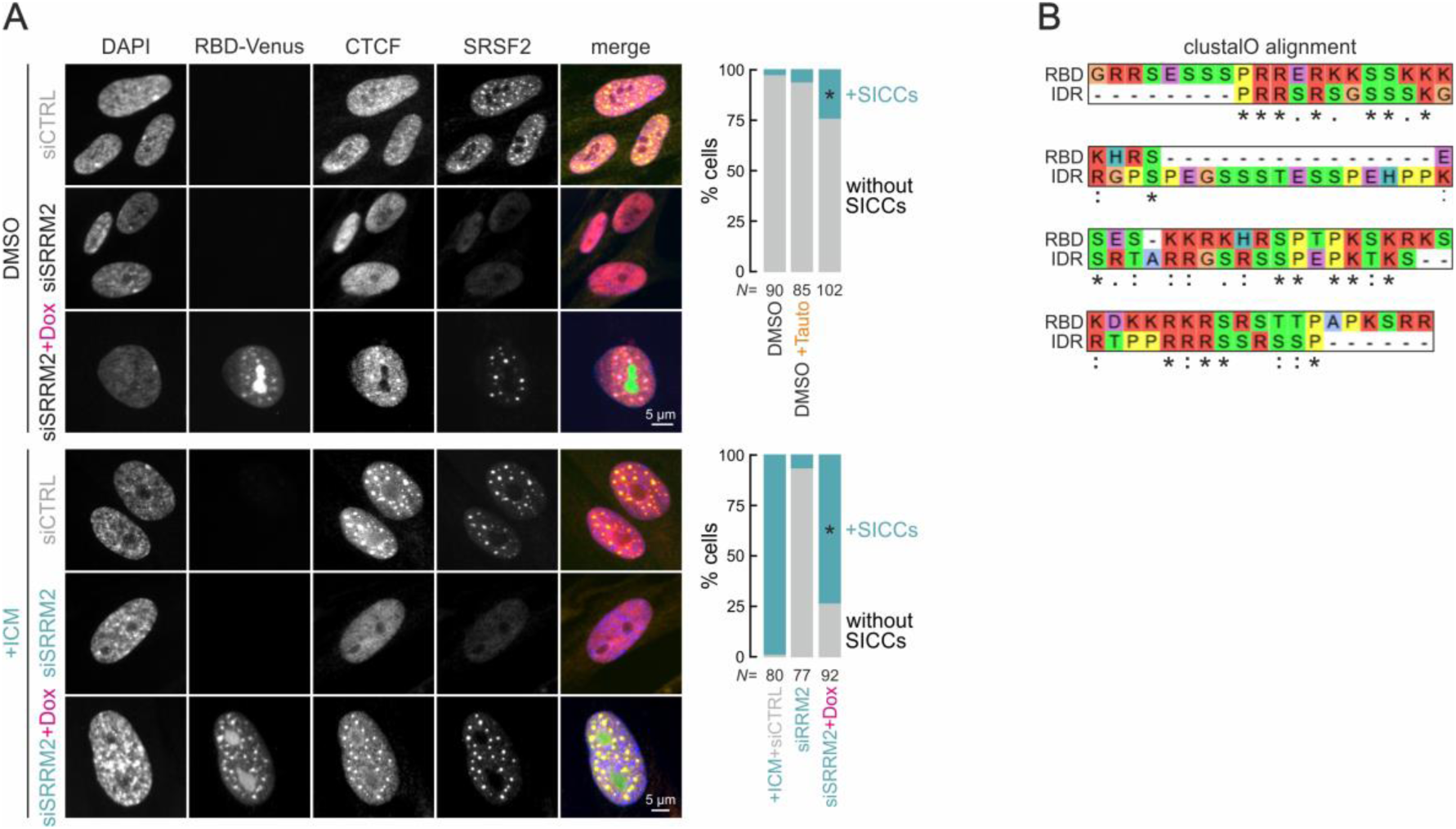
The SRRM2 RNA-binding domain suffices for CTCF clustering (*linked to* Figure 3). **A**, *Left*: Representative immunofluorescence images of proliferating (DMSO) and senescent IMR90 (+ICM) stained for CTCF and SRSF2, and counterstained by DAPI that were treated or not with doxycycline (±Dox) to induce expression of SRRM2 RBD-Venus fusion constructs (green channel) in cells carrying or not full-length SRRM2 because of its siRNA-mediated knockdown. *Right*: Bar plots showing the percentage of cells displaying SICCs; *N* is the number of cells quantified in each condition. \**P*<0.05, Fisher’s exact test. **B**, Alignment of the 68 aa-long sequences of the SRRM2 RNA-binding (RBD) and its C-terminal IDR domains (IDR; see Figure 3E). Asterisk = conserved amino acid residue; colon = highly similar aa residue properties (>0.5 in the Gonnet PMA matrix); period = weakly similar aa residue properties (<0.5 in the Gonnet PMA matrix).

**Figure S8.**
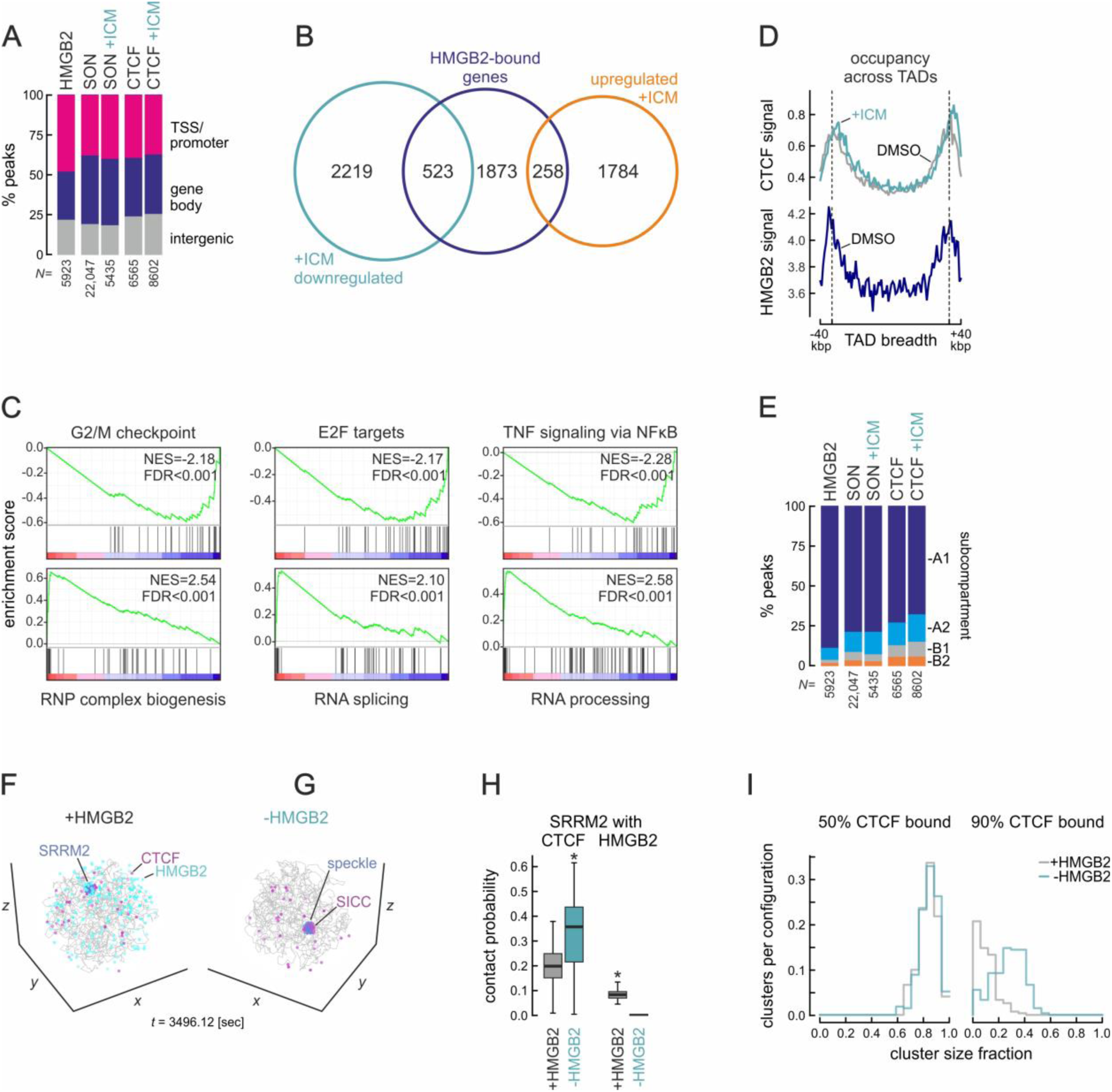
HMGB2 in 3D chromatin organization and senescence entry (*linked to* Figure 4). **A,** Bar plots showing the genomic distribution of HMGB2 and SON ChIPmentation and of CTCF CUT&Tag peaks in proliferating (DMSO) and senescent IMR90 (+ICM). *N* is the number of peaks annotated in each condition. **B,** Venn diagram showing the overlap of HMGB2-bound genes (blue) with genes differentially up-(orange) or downregulated upon ICM treatment (green). **C,** GSEA analysis of the shared HMGB2-bound and up-/downregulated genes from panel B. **D,** Plots showing scaled CTCF (top) and HMGB2 binding signal distribution relative to all TADs from Micro-C data ±40 kbp up-/downstream of TAD boundaries. **E,** Bar plots showing the subcompartmental distribution of the HMGB2, SON, and CTCF peaks from panel A. *N*, the number of peaks annotated in each condition. **F,** 3D renderings of the near-equilibrium MD models with (left) or without HMGB2 (right) used to assess SICC formation relative to phase-separating speckles of SRRM2. **G,** Box plots showing the contact probability of SRRM2 with CTCF or HMGB2 beads in MD simulations. \**P*<0.01, unpaired two-tailed Student’s t-test. **H,** Plots showing the number of CTCF clusters and their sizes in MD simulations with (grey line) and without HMGB2 (green line) given increasing fractions of chromatin-bound CTCF.

**Figure S9.**
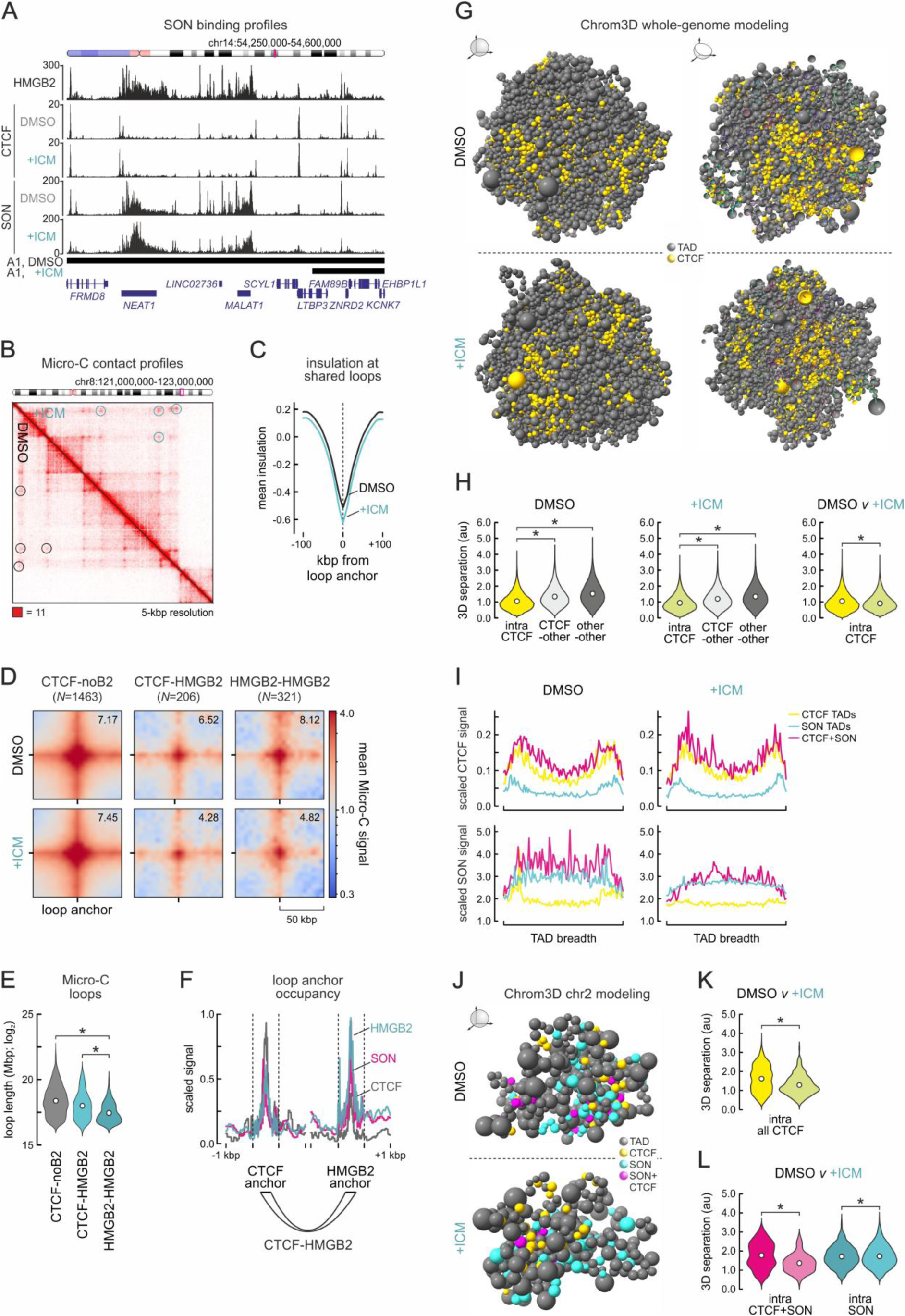
Micro-C data and Chrom3D models of 3D chromatin reorganization in senescence (*linked to* Figure 5). **A**, Representative genome browser view of HMGB2 and SON ChIPmentation data aligned to CTCF CUT&Tag profiles from proliferating (DMSO) and senescent IMR90 (+ICM) in a 350-kbp region on chr14. **B**, Contact matrix showing Micro-C interactions from proliferating (DMSO) and senescent IMR90 (+ICM) in a 2-Mbp region on chr8. CTCF loops are indicated (black/green circles). **C**, Plots showing mean insulation scores from proliferating (DMSO) and senescent Micro-C (+ICM) in the 200 kbp around loop anchors shared in between the two conditions. **D**, APA plots showing mean Micro-C signal strength in proliferating (DMSO) and senescent cells (+ICM) for loops with CTCF-but no HMGB2-bound anchor, with one CTCF- and one HMGB2-bound anchor or with two HMGB2-bound anchors in proliferating cells. **E**, Violin plots showing length distribution of the three loop groups from panel D. \**P*<0.01; Wilcoxon-Mann-Whitney test. **F**, Line plots showing CTCF (grey), HMGB2 (green) and SON occupancy (magenta) in the 2 kbp around the anchors of loops with CTCF on one side and HMGB2 in the other. **G**, Renderings of Chrom3D diploid genome models from proliferating (DMSO) and senescent IMR90 (+ICM) showing TADs (beads) that carry the top 20% of CTCF signal (yellow). **H**, Violin plots showing mean pairwise 3D separation of the indicated types of beads in each model from panel D. \**P*<0.01; Wilcoxon-Mann-Whitney test. **I**, Plots showing scaled CTCF (top) or SON occupancy (bottom) within CTCF-(yellow), SON-(light blue) and CTCF+SON-enriched TADs (magenta) from the proliferating (DMSO) and senescent Chrom3D models (+ICM). J, As in panel G, but from chr2 alone and indicating CTCF-(yellow), SON-(light blue) and CTCF+SON-enriched TADs (magenta). **K**, As in panel H, but only using 3D separation values from the models in panel G. **L**, As in panel K, but assessing separation among SON-(light blue) and CTCF+SON-enriched TADs (magenta).

**Figure S10.**
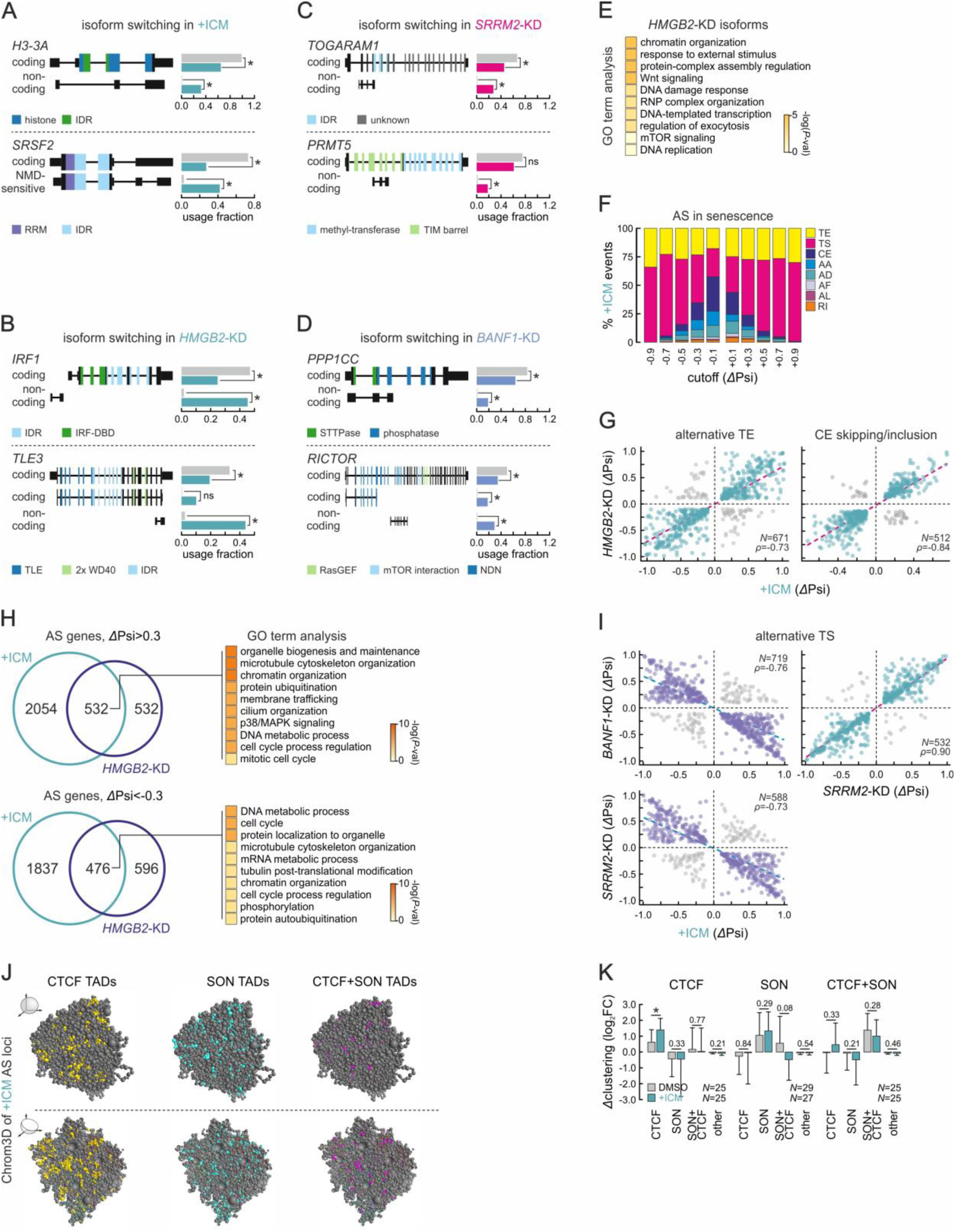
Alternative splicing patterns upon senescence entry (*linked to* Figure 6). **A**, Exemplary changes in mRNA isoforms deduced from the analysis of mRNA-seq data from senescent IMR90 (+ICM). \**P*<0.05, unpaired two-tailed Student’s t-test. **B**, As in panel A, but from *HMGB2*-knockdown data. **C**, As in panel A, but from *SRRM2*-knockdown data. D, As in panel A, but from *BANF1*-knockdown data **E**, Heat map of the top 10 GO terms/pathways enriched for AS isoforms emerging upon *HMGB2*-KD. **F**, Bar plots showing the percent of AS event types at different *Δ*Psi cutoffs in senescence (+ICM) RNA-seq data. **G**, Scatter plots showing correlation of *Δ*Psi values of alternative transcription-end sites (TE) shared between senescence (+ICM) and *HMGB2*-KD. The number of TE events (*N*) and correlation coefficients (ρ) are indicated. H, Venn diagrams showing the overlap of AS genes with a *Δ*Psi>0.3 (top) or <-0.3 (bottom) between senescence (+ICM) and *HMGB2*-KD. The top 10 GO terms/pathways enriched for shared genes are shown (right). **I**, As in panel G, but for alternative transcription-start site (TS) events shared between senescence (+ICM) and *BANF1*- or *SRRM2*-KD. **J**, Renderings of Chrom3D diploid genome models from senescent IMR90 showing TADs (beads) that carry the top 20% of CTCF (yellow), SON (light blue), or CTCF+SON signal (magenta) and AS genes from +ICM data. **K**, Bar plots showing clustering (±SEM) of CTCF (left), SON (middle) or CTCF+SON TADs (right) from panel J relative to randomized controls in DMSO-(grey) or ICM-treated cells (green). \**P*<0.05; unpaired two-tailed Student’s t-test. The number (*N*) of clusters deduced in each case is indicated.

**Figure S11.**
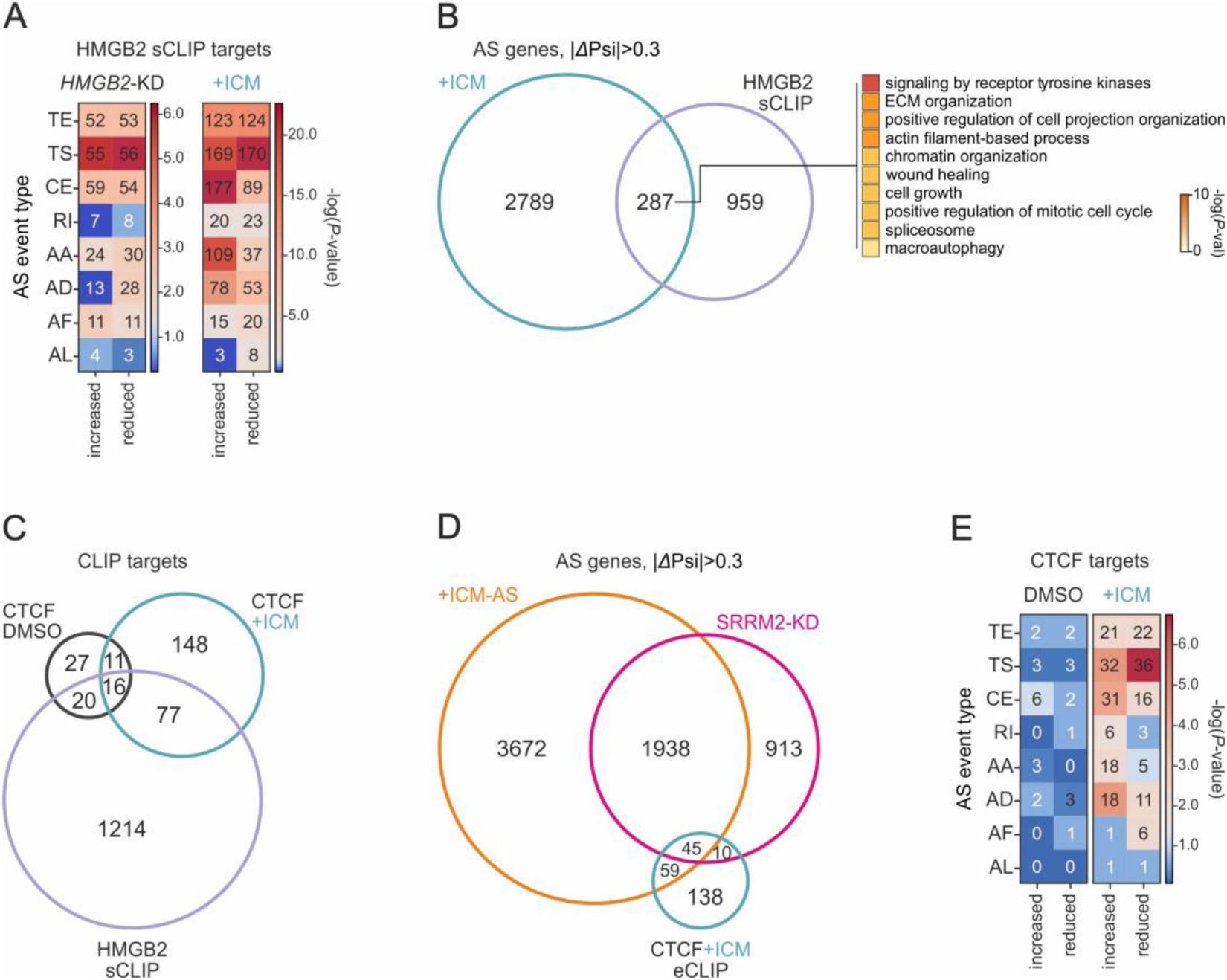
CLIP targets of HMGB2 and CTCF that are alternatively spliced in senescence (*linked to* Figure 6). **A**, Heat map showing the number and relative enrichment [-log(*P*-value)] of AS events from HMGB2 target-mRNAs in *HMGB2*-KD and senescent IMR90 (+ICM). **B**, *Left*: Venn diagram showing the overlap of AS genes in senescence (+ICM) with HMGB2 sCLIP targets in proliferating IMR90. *Right*: The top 10 GO terms/pathways enriched for the 287 shared targets. **C**, Venn diagram showing the overlap of CTCF- and HMGB2-bound mRNAs from CLIP data. **D**, As in panel C, but for the overlap of AS genes in senescence (+ICM) or SRRM2-knockdown (KD) with CTCF eCLIP targets in ICM-treated IMR90. **E**, As in panel A, but for CTCF-bound mRNAs in proliferating (DMSO) and senescent IMR90 (+ICM).

Table S1. CTCF interactome profiling via co-IP MS/MS (provided as an .xlsx file).

Table S2. Analysis of alternative splicing patterns via IsoformSwitch (provided as an .xlsx file).

**Table S4.**
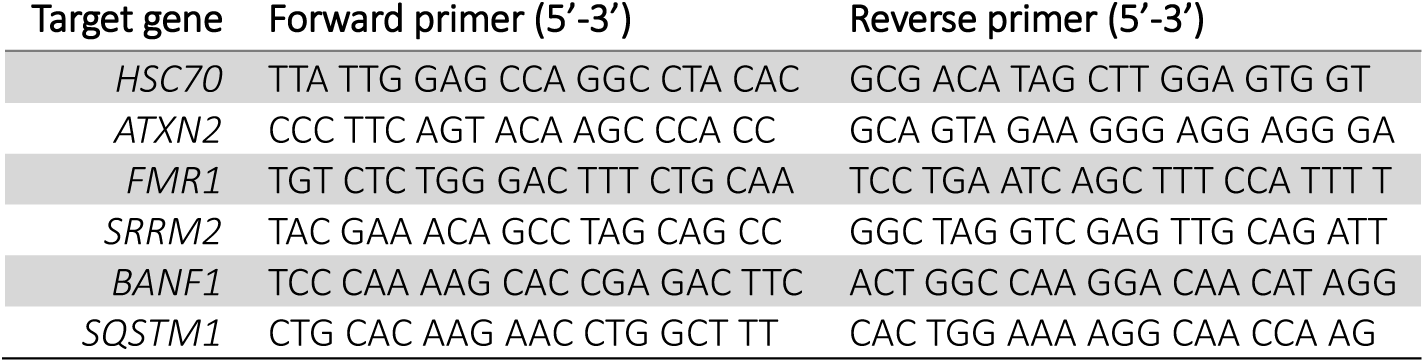
List of qPCR primers used in this study.

**Table S5.**
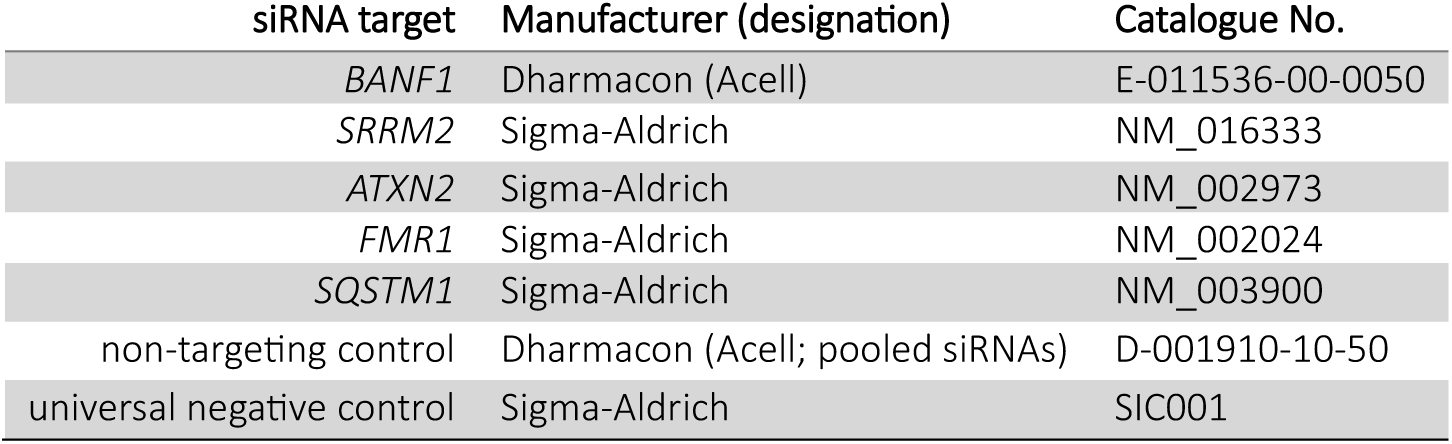
List of siRNAs used in this study.

| siRNA target | Manufacturer (designation) | Catalogue No. |
| --- | --- | --- |
| *BANF1* | Dharmacon (Acell) | E-011536-00-0050 |
| *SRRM2* | Sigma-Aldrich | NM_016333 |
| *ATXN2* | Sigma-Aldrich | NM_002973 |
| *FMR1* | Sigma-Aldrich | NM_002024 |
| *SQSTM1* | Sigma-Aldrich | NM_003900 |
| non-targeting control | Dharmacon (Acell; pooled siRNAs) | D-001910-10-50 |
| universal negative control | Sigma-Aldrich | SIC001 |

Table S3. Analysis of alternative splicing patterns via Whippet (provided as an .xlsx file).

Table S6. Catalogue of ChIPmentation peaks (provided as an .xlsx file).

Table S7. Catalogue of HMGB2 and CTCF CLIP targets (provided as an .xlsx file).

